# Local GEBVs and additive variance in maize heterotic groups across low- and high-stress environments

**DOI:** 10.64898/2026.11.06.763794

**Authors:** Marcos Antonio de Godoy Filho, Caio Canella Vieira, Diego Jarquín, Kaio Olímpio das Graças Dias, José Baldin Pinheiro

## Abstract

Hybrid maize (*Zea mays* L.) parents are chosen within heterotic groups. Drought and heat stress are expected to become more frequent, but little is known about how they change the additive variance of each group and where in the genome it lies. We analyzed testcross grain yield from 62 Genomes to Fields environments, classified as low or high stress by climate. For 1,362 parents from three heterotic groups and two doubled-haploid families, we fitted a genomic model with group deviations and calculated local genomic estimated breeding values in 164-kb segments. In the heterotic groups, high stress reduced additive variance by 40–57%, and the genetic correlation between conditions was 0.73–0.99. Within groups, the distribution of variance across the genome was similar between conditions (correlation 0.84–1.00), and genotype-by-environment interaction was spread over many low-variance segments. Genome-wide association detected 25 loci under low stress and 18 under high stress, none shared, but half or more lay in high-effect, high-variance segments that were nearly the same in both conditions. The best possible cross between the two heterotic pools exceeded the best evaluated hybrid by 105–117%, a gap that depended on segment size, not on stress. With 25 founders, stacking the best segments reached 94–95% of the pool potential, versus 56–92% for selection on total breeding value, whose founders were more related. Founders chosen in one condition reached, in the other, 94–100% of the potential of those chosen there. Local breeding values can guide the choice of donors and founders under both conditions.

**Article summary:** Hybrid maize breeders choose parents within heterotic groups, and drought and heat stress are expected to increase. Using public Genomes to Fields trials, we measured how stress changes the additive genetic variance of each group and where it lies in the genome. Stress reduced the variance in the three heterotic groups, but the regions with the most variance were nearly the same under low and high stress. Founders chosen by combining the best genome segments reached almost all of the pool’s potential and kept it under the other condition. Breeders can use segment values to choose parents across stress levels.

## Introduction

Genomic selection (GS) predicts the additive genetic value of each individual as the genomic estimated breeding value (GEBV), based on markers distributed across the whole genome (Meuwissen et al. 2001). In hybrid maize, GS was preceded by the prediction of single-cross hybrids from related hybrids (Bernardo 1994, 1996). Today, it is routinely used in plant breeding (Crossa et al. 2017), including large commercial maize breeding programs (Cooper et al. 2016). In their most widely used form, the models treat the panel as a single population and predict mean performance in the training environments. However, because of genotype × environment interaction (G×E), the parent or hybrid with the best mean may not be the best in contrasting conditions, such as water and heat stress, which is expected to become more frequent in the Corn Belt with climate change (Ting et al. 2021). For years and regions with stress, breeders need to choose parents, not only to predict performance. Selection on GEBV chooses the individuals with the highest total value, but the best version of each stretch of the genome is spread across the population, including individuals that are not among those with the highest GEBV. In wheat, chickpea and dairy cattle, combining these versions produced predicted values above the best individual (Hayes et al. 2024).

To account for G×E, prediction began to use reaction norms with environmental covariates (Jarquín et al. 2014) and crop growth models integrated with genomic prediction, developed in commercial maize breeding (Cooper et al. 2016; Messina et al. 2018). The Genomes to Fields (G2F) project has evaluated hybrids in North America since 2014, with publicly available genotypes and climate data (AlKhalifah et al. 2018; McFarland et al. 2020). It brings together hundreds of environments for the study of G×E (Washburn et al. 2025). Most of the hybrids are testcrosses of lines with a small number of testers (Lima et al. 2023; Lopez-Cruz et al. 2023). These studies address the prediction of performance. Little is known about how stress changes the additive variance available for selection and its distribution across the genome.

In the United States, maize hybrids are crosses between a small number of heterotic groups with a narrow genetic base (Duvick 2005; Mikel and Dudley 2006), and each group is improved separately (Hallauer et al. 2010). Therefore, parents are chosen within each group, and the response to selection depends on the variance of the group (Silva et al. 2020). The groups differ in marker effects (Technow et al. 2014) and in genetic variance. In G2F, the non-Stiff Stalk group had a larger general combining ability variance than the Stiff Stalk group (Godoy dos Santos et al. 2026). Under stress, the response to selection depends on the variance and heritability in that condition. When selection is carried out under low stress, the response under stress also depends on the genetic correlation between conditions (Rosielle and Hamblin 1981). When this correlation is below 1, the ranking of genotypes changes between conditions, which affects selection (Cooper and Messina 2023). If stress reduces the variance of one group more than that of another, the response to selection under stress differs among groups. The choice of parents should also consider how variance is distributed across the genome.

The genome can be divided into segments, contiguous stretches of a chromosome, and the GEBV of each parent can be decomposed into the sum of its values in the segments. The value of a parent in a segment is the local genomic estimated breeding value (local GEBV; Voss-Fels et al. 2019). With the local GEBV, the choice of parents does not depend only on total GEBV. Stacking combines the best segment of each part of the genome into a single genotype, and the value of this genotype (the stacking ceiling) is a selection limit (Cole and VanRaden 2011; Kemper et al. 2012). Stacking is also used to choose founders, parents that together carry the best segments. In simulations, stacking maintained more diversity and had greater long-term gain than truncation, which is the choice of the parents with the highest GEBV (Villiers et al. 2024; Shaffer et al. 2026a). Founders can form the population of a GS program (Kemper et al. 2012; Villiers et al. 2024; Shaffer et al. 2026a) or bring new germplasm into an ongoing breeding program, such as the donors chosen from maize testcross data to enrich elite lines (Allier et al. 2020).

The ceiling depends on segment size, from the whole chromosome to the single marker (Cole and VanRaden 2011; Hayes et al. 2024). In maize, marker effects were used in simulations to identify the recombination points that would maximize gain on each chromosome of a biparental cross (Bernardo 2017). In hybrid maize, the product is the cross between heterotic pools, and the ceiling is that of the cross between the best possible lines of the two pools. In wheat, stacking was used to improve the stability of disease resistance across environments (Tong et al. 2025). If segment variance changes with stress, the best segments and the chosen founders may also change. Local GEBVs usually come from models in which all markers have the same variance, and the effect of a locus is spread over the segment. A genome-wide association study (GWAS) tests one marker at a time, and the effect of climate can be studied locus by locus. In maize testcross hybrids, almost all loci for grain yield had effects that depended on the environmental scenario, and those for heat and drought tolerance had strong effects only in hot or dry scenarios (Millet et al. 2016). In G2F, the quantitative trait loci (QTL) for grain yield showed high variation across environments and testers (Ozair et al. 2025). Each locus lies in a segment whose variance can be measured with local GEBVs, and, in wheat, the highest-variance blocks were compared with known QTL (Tong et al. 2025).

In this study, we applied to G2F hybrid maize a GWAS and the segment-based analyses used in stacking, with the population divided into heterotic groups and the environments into two conditions. The rainfed environments in the northern United States and Canada were divided into low and high stress based on evaporation at flowering and temperature during vegetative growth. The parents of the hybrids form three heterotic groups (the Stiff Stalk, non-Stiff Stalk and Iodent groups) and two doubled-haploid (DH) families. We fitted a genomic best linear unbiased prediction (GBLUP) model with group deviations, in which condition was a fixed effect and the genetic effect of each parent had a variance in each condition and a covariance between conditions. We then calculated the local GEBV of each parent in each segment of the genome, as in stacking.

The general objective was to evaluate how stress changes the additive variance of each heterotic group, in total and across the genome, and what this implies for the choice of parents and founders by local GEBV. The specific objectives were to measure how much stress changes the additive variance and the genetic correlation of each group and whether it changes the distribution of variance across the genome or only its level, and to estimate the stacking ceiling between the two heterotic pools, from the whole chromosome to the single marker, in both conditions. Other objectives were to compare the choice of founders by stacking and by truncation, to measure how much of their potential the founders chosen in one condition retain in the other and to place the GWAS loci relative to segment variance. Because each target region includes low- and high-stress years, and the proportion between them changes with climate (Cooper and Messina 2023), we aimed to provide breeders with information to choose parents and segment donors in ongoing breeding programs, and founders when a program starts or is renewed, for low- and high-stress years and regions.

## Materials and methods

### 2.1 Panel, genotypes and relationship matrix

We used the hybrid trials of the Genomes to Fields (G2F) project from 2014 to 2023. The hybrids are testcrosses between inbred lines and a small set of testers. Twelve testers appear in 30 or more of the analyzed hybrids and account for most of the panel: LH195, LH244, PHT69, PHZ51, PHP02, PHB47, LH185, LH198, LH82, CG102, DK3IIH6 and PB80.

The genotypes of the parents were obtained from the G2F 437,214-marker set for inbred lines (Practical Haplotype Graph, B73 v5 coordinates; Washburn et al. 2025). We excluded parents with heterozygosity above 5% and kept polymorphic markers with minor allele frequency (MAF) ≥ 0.05 and less than 10% missing data. This left 2,075 parents and 333,382 markers. Missing data were imputed with the markov function of bWGR 2.2.13 (Xavier et al. 2020).

The genomic relationship matrix followed VanRaden (2008): **G** = **MM**^⊤^/*c*, where **M** is the matrix of centered genotypes and *c* = 2 ∑ *p*(1 − *p*).

### 2.2 Trials and first stage

We restricted the network to the northern United States and Canada (latitude ≥ 37° N), to trials up to 2023 and to environments (location × year) with at least 15 hybrids. This left 192 environments, 4,966 hybrids and 78,485 observations.

The first stage followed Godoy Filho et al. (2026a). An environment was defined as a unique combination of location and year. Therefore, different trials conducted at the same location were analyzed together, as a single environment, because most were planted on similar dates. Most trials followed randomized complete block designs or incomplete block designs, generally with two replicates. The analysis was performed in two stages, and environments were first analyzed individually to obtain adjusted means, the best linear unbiased estimates (BLUEs). To accommodate the diversity of designs and the presence of environments with more than one trial, the model structure was adjusted dynamically, environment by environment, in ASReml-R 4.2 (Butler et al. 2023). The general linear mixed model was:

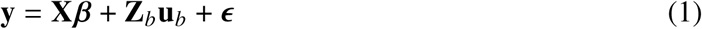

where **y** is the *n* × 1 vector of observed phenotypic values (grain yield), with *n* the total number of plots in the environment; ***β*** is the *p* × 1 vector of fixed effects, which includes the intercept, genotype, replicate and trial (when applicable); **X** is the *n* × *p* incidence matrix that links the observations to the fixed effects; **u***_b_* is the *q* × 1 vector of random block effects, nested within trials and replicates in incomplete block designs, with 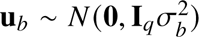, where **I***_q_* is the identity matrix of order *q*; and **Z***_b_* is the *n* × *q* incidence matrix of the random block effects. To account for the diversity of designs and possible heteroscedasticity, the model structure was adjusted dynamically. In environments with more than one trial, the fixed effect of trial was included and a heterogeneous residual variance structure was modeled, ***ε*** ∼ *N* (**0**, **R**), with 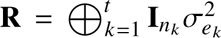, where *t* is the number of trials and *n_k_* is the number of plots in the *k*th trial. To assess trial quality, generalized heritability 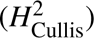 was estimated following Cullis et al. (2006). For this, the models were refitted with genotype effects as random, 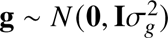. Generalized heritability was calculated as:

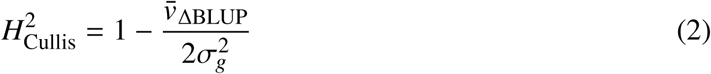

where *v̄*_ΔBLUP_ is the average pairwise prediction error variance of the best linear unbiased predictions (BLUPs) and 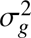 is the genotypic variance. The coefficient of variation of the adjusted means of each environment was calculated as:

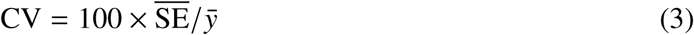

where SE̅ is the mean standard error (SE) of the BLUEs of the environment and *ȳ* is the mean of the BLUEs. We excluded irrigated trials and environments with 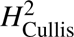 < 0.10 or CV above 25%. This left 147 environments.

### 2.3 Climate covariates and phenological windows

Climate covariates were obtained from the NASA Prediction Of Worldwide Energy Resources project (NASA POWER) with nasapower (Sparks 2018) and processed with EnvRtype (Costa-Neto et al. 2021) in phenological windows (Godoy Filho et al. 2026b). A phenological window is an interval of the cycle defined in growing degree days before and after flowering. Thus, each window corresponds to the same phase of development in all environments. We used seven windows: W1, 0–40% of the growing degree days from planting to flowering; W2, 40–90%; W3, 90–100%; W4, from flowering to +10% of the growing degree days of the reproductive phase; W5, +10–30%; W6, +30–65%; W7, +65–100%. The value of a covariate in a window is the sum of the daily values in the window. Each of the 33 variables was used in nine forms: in the seven windows, as the mean over the seven windows and as the cycle total (the sum of the seven windows), for a total of 297 covariates.

### 2.4 Low- and high-stress conditions

To define the stress conditions, we used an environmental index, a measure of the yield level of each environment: the mean of the stage 1 BLUEs of the 20 hybrids of that environment that were evaluated in the largest number of environments of the network. The index is based on yield only, and the climate covariates enter afterwards, in the choice of the grid.

The two conditions were defined by a grid (Figure 1a). A grid divides the environments into 2 × 2, 2 × 3, 3 × 2 or 3 × 3 cells by splitting two covariates at the median or at the terciles. First, among the 297 covariates, we preselected the 90 (30%) with the highest absolute correlation with the environmental index. Then, we tested all pairs of these covariates derived from different variables and weakly correlated with each other (|*r* | < 0.50), in the four grid layouts. We required at least 25 environments in each extreme cell. The criterion was the largest difference in the environmental index between the highest- and the lowest-yielding cells, measured by the lower bound of the 95% confidence interval (95% CI) of that difference (t-test).

**Figure 1.**
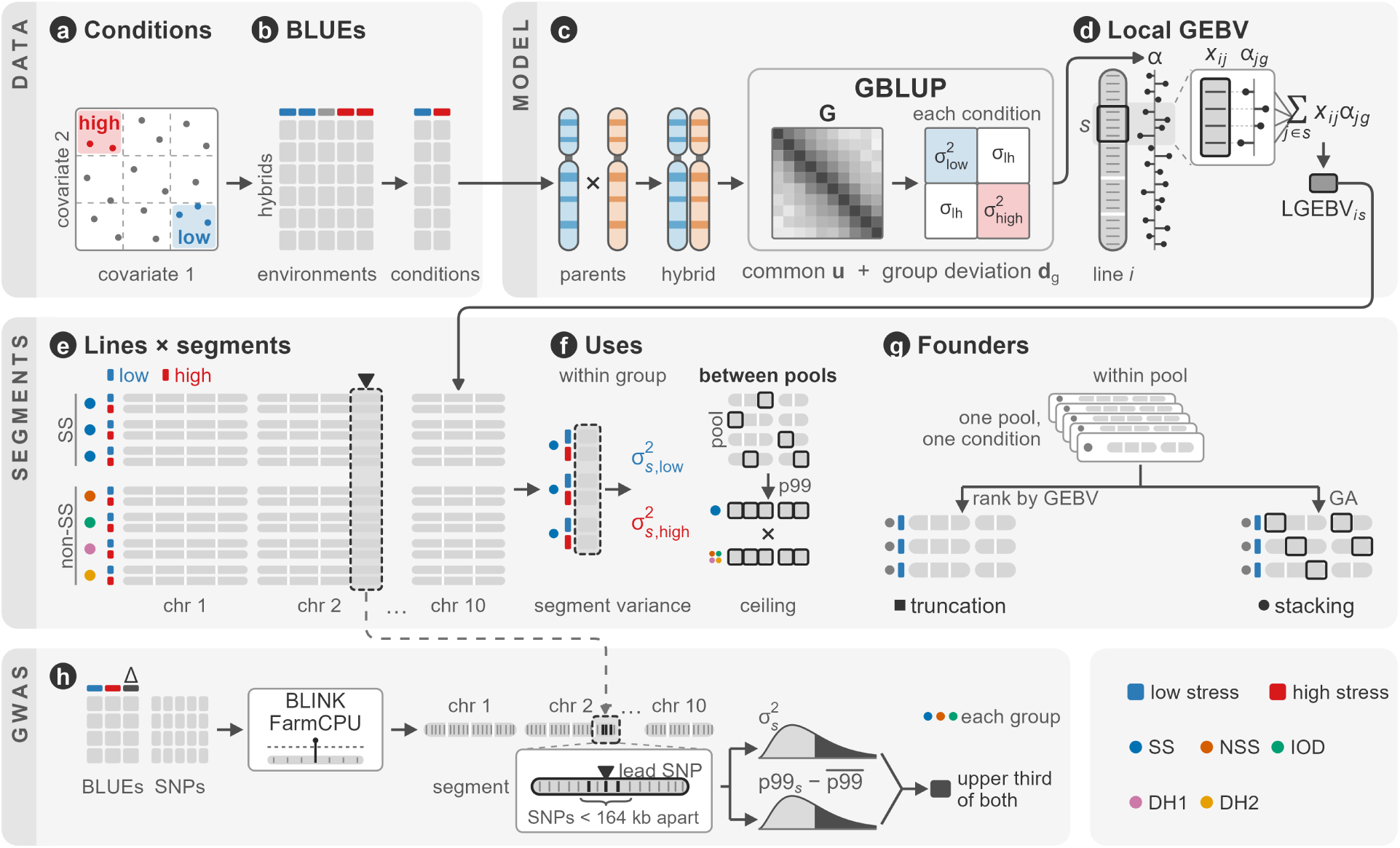
Outline of the methods. (a) The environments were divided into low and high stress by a grid of two climate covariates. (b) The hybrid BLUEs were obtained in each environment (stage 1) and then in each condition (stage 2). (c) The GBLUP decomposes the genetic effect of each parent into a common effect (**u**) and its group deviation (**d***_g_*), with a variance in each condition and a covariance between conditions. (d) The marker effects (*α*) come from the GEBVs. The local GEBV of line *i* in segment *s* is the sum, over the markers in the segment, of the genotype times the marker effect. (e) In each condition, the local GEBVs form a matrix of lines by segments, with the parents of the Stiff Stalk and non-Stiff Stalk pools in separate sets of rows. (f) Within each group, the segment variance in each condition. Between the pools, the stacking ceiling: the 99th percentile of each segment forms the best possible line of each pool, and the two are crossed. (g) Within each pool and condition, the founders are chosen by total GEBV (truncation) or by the genetic algorithm (stacking). (h) GWAS with BLINK and FarmCPU on the low-stress BLUEs, the high-stress BLUEs and the delta (Δ). Each locus groups the associated SNPs less than 164 kb apart. The segment of the lead SNP is compared, in each group and condition, with the distribution of the variance 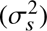 and that of the 99th percentile of the local GEBV of the segment minus the mean of the 99th percentiles of all segments (p99*_s_* – mean). Segments in the upper third of both are the high-effect, high-variance segments. Colors identify the conditions and the groups: Stiff Stalk (SS), non-Stiff Stalk (NSS or non-SS), Iodent (IOD) and the DH families PHN11 × PHW65 (DH1) and PHW65 × Mo44/MoG (DH2). GA, genetic algorithm; LGEBV, local GEBV.

The highest-yielding cell formed the low-stress condition, and the lowest-yielding cell formed the high-stress condition. Environments in the other cells were not used in the analyses by condition. We also excluded environments with less than 5% hybrid connectedness with the other environments of the same condition. This left 62 environments, 28 under low stress and 34 under high stress (Supplementary Table S1).

### 2.5 Second stage

In stage 2, we fitted the following model for each stress condition with ASReml-R:

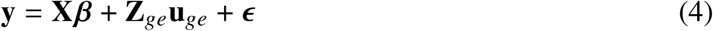

where **y** is the vector of stage 1 BLUEs in the environments of the condition; ***β*** is the vector of fixed environment and hybrid effects; **u***_ge_* is the vector of random hybrid × environment interaction effects, with 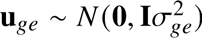; **X** and **Z**_*ge*_ are the incidence matrices; and 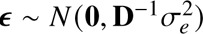, where **D** is the diagonal matrix of the weights 1/SE^2^ from stage 1. The result was one BLUE per hybrid per condition, with its SE (Figure 1b). In the following analyses, we used only the hybrids whose two parents have hybrids evaluated in both conditions.

### 2.6 Germplasm groups

We clustered the 2,075 parents with k-means (k = 5, 500 random starts). Each group was named after the heterotic group of its predominant individuals. The three heterotic groups were the Stiff Stalk group (770 parents), the non-Stiff Stalk group (Lancaster/Oh43 lines and 394 GEM-derived lines, Germplasm Enhancement of Maize, 680 parents) and the Iodent group (119). The other two were DH families from the G2F biparental populations with PHW65 as the common parent, PHN11 × PHW65 (192) and PHW65 × Mo44/MoG (314) (Lima et al. 2023; Costa et al. 2024). The group of each parent is given in Supplementary File S1.

### 2.7 Genetic model with group deviations

We fitted a GBLUP to the stage 2 BLUEs with ASReml-R (Figure 1c). There were 5,119 BLUEs of 3,102 hybrids, from 1,362 parents with hybrids evaluated in both conditions. As in de los Campos et al. (2015) and Rio et al. (2020), the model decomposes the genetic effect of each parent into an effect common to all parents and a deviation specific to its group. Condition was fitted as a fixed effect, and each of the two parts had a variance in each condition and a covariance between conditions:

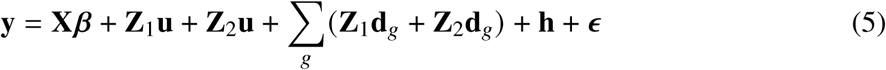

where **y** is the vector of stage 2 BLUEs; ***β*** is the vector of condition means, and **X** is its incidence matrix; **u** is the vector of common genetic effects of the parents, with incidence matrices **Z**_1_ and **Z**_2_ for the first and second parent of each hybrid, and **u** ∼ *N* (**0**, **G** ⊗ **Σ***_u_*), where **Σ***_u_* is the 2 × 2 unstructured matrix of the genetic variances in the two conditions and the covariance between them, and ⊗ is the Kronecker product; **d***_g_* is the vector of deviations of group *g* (the five groups of section 2.6), with **d***_g_* ∼ *N* (**0**, **G***_g_* ⊗ **Σ***_g_*), where **G***_g_* is **G** restricted to the group (the submatrix of the members of *g* and 10^−6^ on the diagonal for the other parents) and **Σ***_g_* is also 2 × 2 unstructured; **h** is the vector of hybrid effects in each condition, which contains the specific combining ability, with **h** ∼ *N* (**0**, **I** ⊗ **Σ**_ℎ_), where **Σ**_ℎ_ is diagonal, with one variance per condition; and 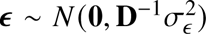, where **D** is the diagonal matrix of the weights 1/SE^2^ from stage 2 and 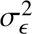 was fixed at 1.

The genetic variance of each group in each condition, which is the testcross additive variance, is the sum of the common variance and the deviation variance, and the genetic correlation between conditions (*r_g_*) was calculated from the variances and covariances. The standard errors were obtained with the delta method (vpredict).

### 2.8 Marker effects and 164-kb segments

Marker effects were back-solved from the GEBVs (Strandén and Garrick 2009): ***α*** = **M**^⊤^**G**^−1^**û**/*c*. The common effect and the deviation of each group were converted into marker effects separately, the common effect with the full **G** and the deviation with the submatrix of **G** for the members of the group. The group-specific marker effect is the sum of the two.

We divided the genome into 10,046 fixed 164-kb segments by physical position. This is the distance at which the mean linkage disequilibrium (LD), measured as *r*^2^ between pairs of markers on the same chromosome, decays to 0.20 in the 4,847 G2F hybrids (Remington et al. 2001). The local GEBV of a parent in a segment is the sum, over the markers in the segment, of the centered genotype times the group-specific effect of the parent’s group (Figure 1d), as in Villiers et al. (2024), Shaffer et al. (2026a) and Tong et al. (2025). The local GEBVs were calculated for the 1,362 parents of the model (section 2.7) and centered on their mean in each segment, so that the sum of the 10,046 local GEBVs of a parent is its GEBV minus the mean GEBV of the 1,362 parents (Figure 1e). The segment variance in a group and a condition is the variance of the local GEBVs among the members of the group (Figure 1f). The marker effects come from an infinitesimal model (GBLUP), with the same variance for all markers, which enter together, without marker selection.

### 2.9 Stacking ceiling and segment resolution

We used two pools of parents, the Stiff Stalk pool (679) and the non-Stiff Stalk pool (non-Stiff Stalk group, Iodent group and the DH families PHN11 × PHW65 and PHW65 × Mo44/MoG, 683 parents). In each pool and condition, the ceiling is the sum, over segments, of the 99th percentile of the local GEBVs among the lines of the pool (Figure 1f). The hybrid ceiling is the ceiling of the Stiff Stalk pool plus that of the non-Stiff Stalk pool, plus the condition mean, in t/ha. The reference is the best evaluated hybrid, the one with the highest predicted value among the hybrids of the model that cross the two pools and were evaluated in the condition. The predicted value is the sum of the GEBVs of the two parents, plus the condition mean. The ceiling assumes that the 99th-percentile local GEBVs of all segments can be combined into a single line, with each segment inherited intact, and does not account for the number of generations required.

To assess how the ceiling depends on resolution, we repeated the calculation with the same marker effects at 13 resolutions by physical position: whole genome, chromosome, one quarter and one fifth of a chromosome, fixed segments of 80, 20, 10, 5 and 1 Mb and of 164 kb, LD blocks (*r*^2^ ≥ 0.5 and ≥ 0.3) and single marker. Several of these resolutions correspond to those used in the literature: the chromosome and the marker, as the lower and upper selection limits of Cole and VanRaden (2011), respectively; segments of about 25 cM, close to one quarter of a chromosome (Kemper et al. 2012); one fifth of a chromosome (Villiers et al. 2024); and LD blocks with *r*^2^ ≥ 0.5 (Tong et al. 2025) and ≥ 0.3 (Shaffer et al. 2026a).

The ceiling was also calculated with the maximum instead of the 99th percentile and with the non-Stiff Stalk pool without the two families (Supplementary Table S2).

To assess how much the ceiling depends on the number of lines, we recalculated the ceiling at 164 kb in the Stiff Stalk and non-Stiff Stalk groups in 30 random samples of 25, 50, 100 and 200 lines. To compare the groups with the same number of lines, we calculated the ceiling of each group (the best possible line of the group) in 50 random samples of 100 lines with phenotypes in the condition.

### 2.10 Founder selection

We used the functions of the HapSelect package (v0.1.0; Shaffer et al. 2026a). In each pool and in each condition, we chose sets of 2, 4, 6, 8, 12, 16, 20, 25 and 50 founders with the local GEBVs of that condition. The sets were chosen by two rules (Figure 1g). In stacking, the set is the one with the highest potential, found by the genetic algorithm of the local_gebv_parent_ selection function, which uses the GA package (Scrucca 2013), with a population of 200 sets in each iteration, up to 400 iterations, and stopping after 80 iterations without improvement. The potential is the sum, over segments, of the mean of the two highest local GEBVs from different founders of the set, in t/ha above the mean of the parents of the model. In truncation, the set consists of the *n* parents with the highest GEBV.

With two founders, the two rules choose the same set.

For each set, we calculated the potential as a fraction of the potential of the whole pool, the mean GEBV, the mean coancestry and the expected heterozygosity. The mean coancestry is the mean of **G**/2 over the pairs of founders. We also measured the diversity per segment, by the number of different versions of each segment among the founders and by the number of founders with the most common version, both averaged over the segments. Two lines have the same version of a segment when they are identical for at least 98% of the single nucleotide polymorphisms (SNPs) of the segment.

Between conditions, we calculated retention, the potential in one condition of the set chosen in the other, divided by the potential of the set chosen in that same condition.

### 2.11 Genome-wide association study (GWAS)

GWAS was performed on the hybrid BLUEs in each condition (section 2.5), using, as in the genetic model (section 2.7), only the hybrids whose two parents have hybrids evaluated in both conditions: 2,756 hybrids under low stress and 2,554 under high stress. We also analyzed the delta, the difference between the two BLUEs (low − high stress), in the 2,084 hybrids evaluated in both conditions. The hybrid genotypes were obtained from the G2F hybrid genotype file, in variant call format. Of the 320,349 markers that had less than 10% missing data, were polymorphic in the hybrids and were present among the 333,382 markers of section 2.1, we retained the 310,948 with MAF ≥ 0.05. Missing data were imputed with the markov function of bWGR 2.2.13 (Xavier et al. 2020).

We used two multi-locus methods from GAPIT 4.1.0 (Lipka et al. 2012; Figure 1h), BLINK (Huang et al. 2019) and FarmCPU (Liu et al. 2016). The covariates were the first three principal components of the hybrid genotypes.

A SNP was considered associated when it reached, with at least one of the two methods, the Bonferroni threshold for the number of independent tests. This number was calculated with simpleM (Gao et al. 2008), which counts, in each LD block, how many eigenvalues of the correlation between genotypes explain 99.5% of the variance and sums these numbers over the blocks. The blocks followed Li et al. (2012): along each chromosome, a SNP joins the block when it has *r*^2^ ≥ 0.1 with any of the 20 preceding SNPs, with up to 1,000 SNPs per block. There were 81,391 independent tests under low stress, 83,111 under high stress and 82,248 for the delta. The threshold, 0.05 divided by this number, was between 6.0 × 10^−7^ and 6.1 × 10^−7^. Associated SNPs less than 164 kb apart (section 2.8) form a locus. The SNP with the lowest *P*-value at each locus (lead SNP) gives the locus its name, with the chromosome and the position in Mb (for example, chr1:7.92). The effect of the locus in each analysis is that of the lead SNP, from the method with the lower *P*-value. The effects refer to the favorable allele, the one that increases grain yield in the condition in which the locus was identified or, for the delta loci, the one that increases the delta. The SNPs significant at a false discovery rate (FDR) of 5% (Benjamini and Hochberg 1995) are listed in Supplementary Table S3. The loci are listed in Supplementary Table S4, with the MAF and the percentage of phenotypic variance explained by the lead SNP (PVE).

To place the loci relative to segment variance, each lead SNP was assigned to the 164-kb segment that contains it (section 2.8). In each segment, we calculated the ratio of the standard deviation of the local GEBV under high stress to that under low stress, among the 1,362 parents of the model, and divided the segments of the genome into thirds by this ratio. The upper third contains the segments with the highest ratio, and the lower third, those with the lowest. In each heterotic group and condition, we also calculated, for each segment, the 99th percentile of the local GEBVs among the parents of the group. From this value, we subtracted the mean of the 99th percentiles of all 10,046 segments. High-effect, high-variance segments are those in the upper third of the distribution of this difference and, at the same time, in the upper third of the distribution of segment variance of the group in the condition. We counted the loci in these segments, in each group, in the condition of the locus.

## Results

### 3.1 Climate contrast and phenotypic data

Of the 5,794 grids tested, the one with the largest yield difference between the extreme cells was the 2 × 3 grid of land-surface evaporation at flowering (W4, median split) and air temperature sum during vegetative growth (W2, terciles; Supplementary Figure S1). Land-surface evaporation in NASA POWER is the evaporation flux over land simulated by the land-surface model of the MERRA-2 reanalysis (Modern-Era Retrospective analysis for Research and Applications, version 2). Low stress corresponded to the cell with high evaporation and the coldest tercile (28 environments), and high stress to the cell with low evaporation and the warmest tercile (34; listed in Supplementary Table S1 and mapped in Supplementary Figure S2). In high-stress environments, the air temperature sum during vegetative growth (W2) was 18% higher (754 vs. 639 °C·day). At flowering (W4), evaporation was 49% lower, evapotranspiration 72% lower, rainfall 51% lower and vapor pressure deficit 82% higher (Figure 2a–e).

**Figure 2.**
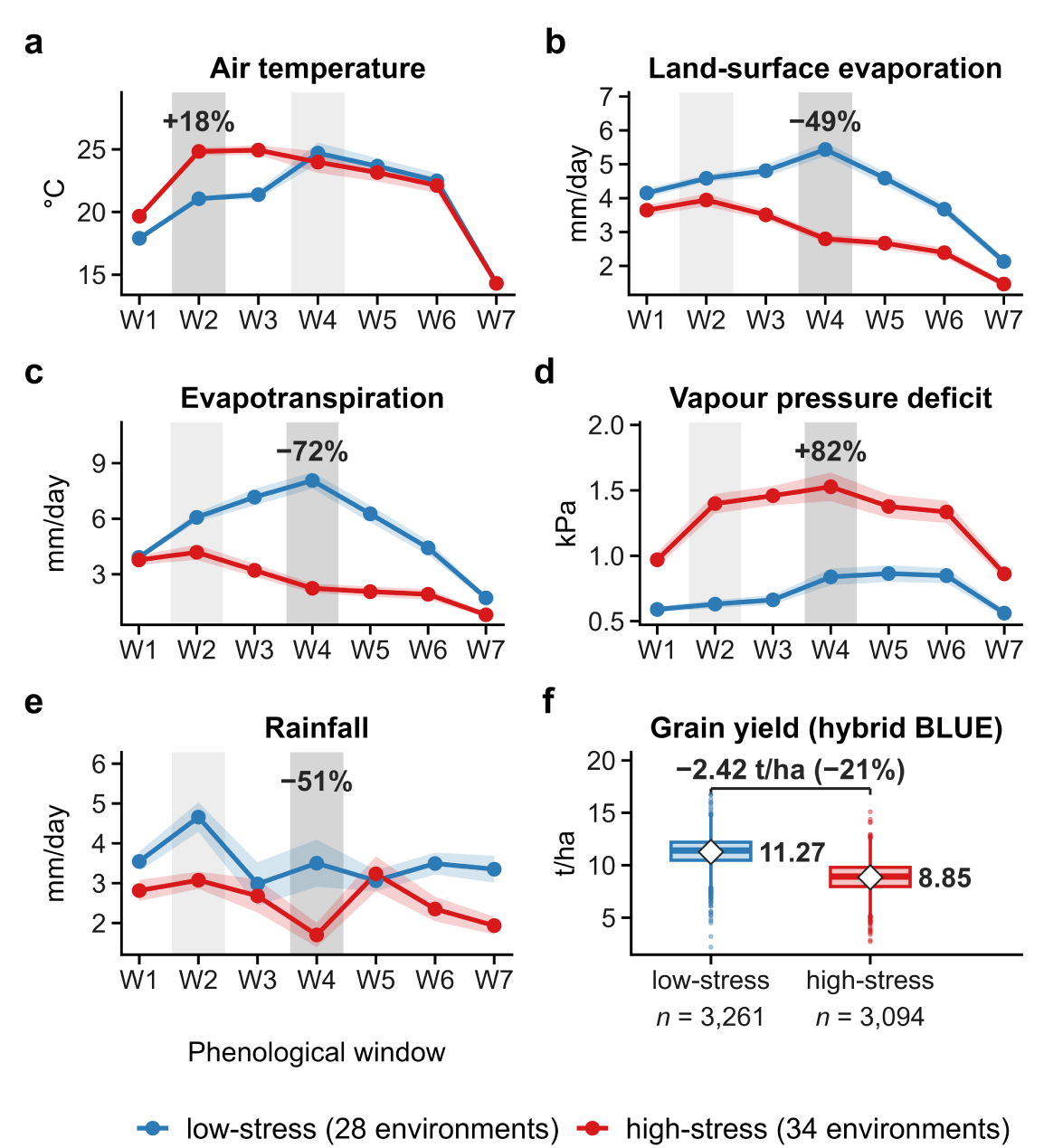
Climate and grain yield in the two conditions, low stress (blue, 28 environments) and high stress (red, 34 environments). Panels a–e show mean air temperature, land-surface evaporation, evapotranspiration, vapor pressure deficit and rainfall in the seven phenological windows (W1–W7), as the window total divided by the mean window length. The lines are the environment means, and the bands show the standard error. The gray background marks the grid windows (W2 and W4), and the value above each panel is the percentage difference of high stress relative to low stress in the darker window. Panel f shows the distribution of hybrid BLUEs, with the mean marked by the diamond and the difference between the means above.

Mean grain yield of the hybrids decreased from 11.27 t/ha under low stress to 8.85 t/ha under high stress (−21%; Figure 2f).

### 3.2 Germplasm structure

The 2,075 parents formed five groups with k-means. Three are heterotic groups, the Stiff Stalk group (770), the non-Stiff Stalk group (Lancaster/Oh43 lines and 394 GEM-derived lines, 680) and the Iodent group (119). The other two groups are DH families from the G2F biparental populations with PHW65 as the common parent, PHN11 × PHW65 (192) and PHW65 × Mo44/MoG (314), which overlap on the first two components and separate on the third (Figure 3a–c). The panel has a single heterotic pattern, because the four groups of the non-Stiff Stalk pool were crossed with the Stiff Stalk group in 83–99.7% of their hybrids (Figure 3d).

**Figure 3.**
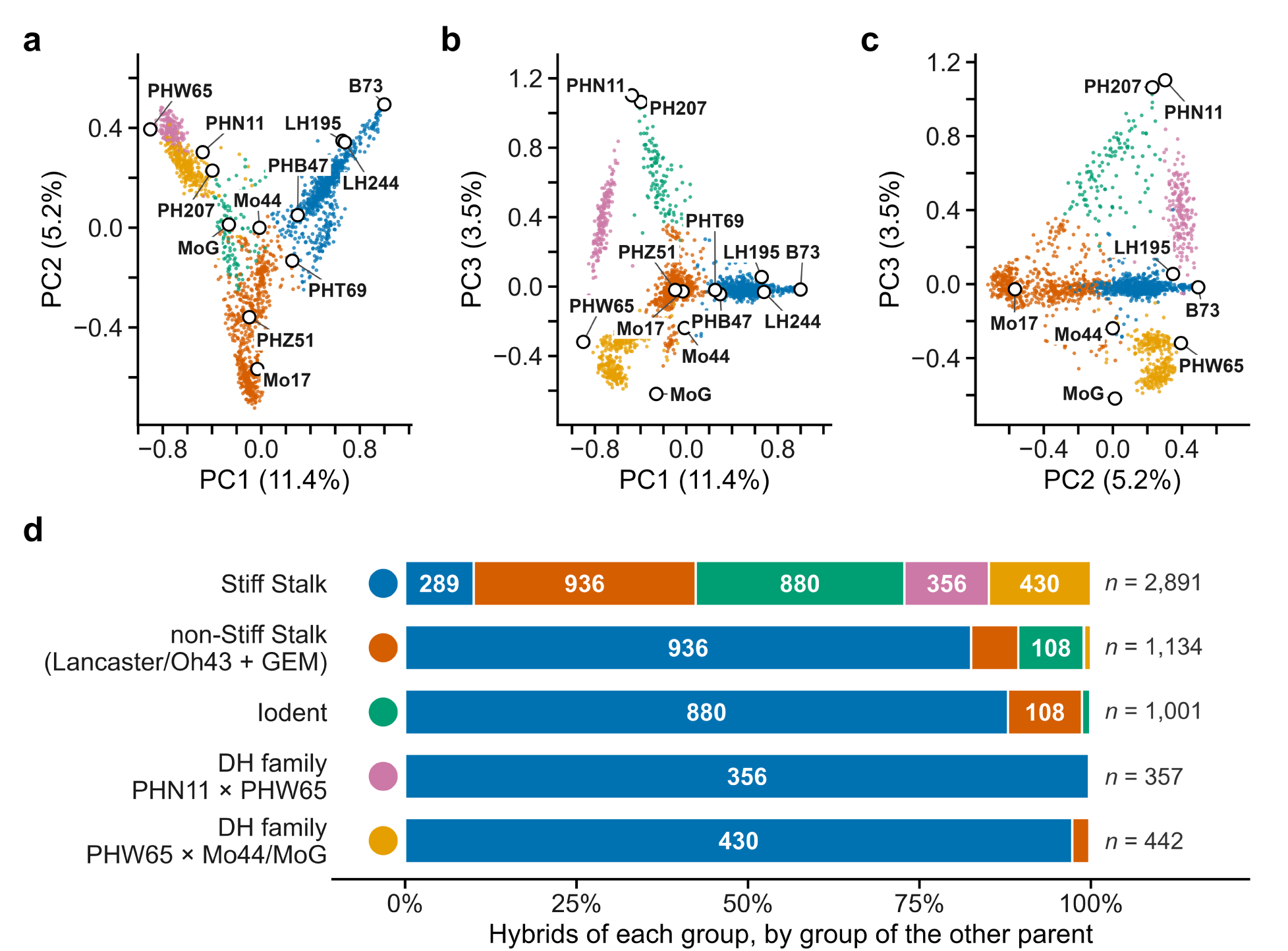
Germplasm structure of the 2,075 parents. Panels a–c show the first three principal components of the genomic relationship matrix, with the percentage of variance on the axes, the five k-means groups in color and the reference lines labeled. Panel d shows, among the hybrids of the genetic model (section 2.7), the percentage of hybrids of each group by group of the other parent, with the number of hybrids on the bars (portions with 7% or more) and the total on the right. The colored circles in panel d identify the groups throughout the figure.

### 3.3 Additive variance by group and condition

The groups differed in variance in both conditions, and high stress reduced the variance of all three. Under low stress, the non-Stiff Stalk group had the largest variance (0.86 ± 0.14 (t/ha)^2^), about twice that of the Stiff Stalk group (0.39 ± 0.05; *P* = 0.001) and that of the Iodent group (0.43 ± 0.10; *P* = 0.013). Under high stress, the non-Stiff Stalk group still had the largest variance (0.41 ± 0.07), above that of the Stiff Stalk group (0.24 ± 0.04; *P* = 0.030) and that of the Iodent group (0.19 ± 0.05; *P* = 0.009). The ratio of high-to low-stress variance was 0.60 in the Stiff Stalk group, 0.47 in the non-Stiff Stalk group and 0.43 in the Iodent group (95% CI below 1 in all three), a decrease of 40%, 53% and 57%, and these percentages did not differ significantly among the groups (*P* ≥ 0.24). In the Stiff Stalk group, the genetic coefficient of variation did not change (5.6% in both conditions). In the non-Stiff Stalk group, the low-stress variance had the widest confidence interval among the groups (Table 1).

**Table 1.**
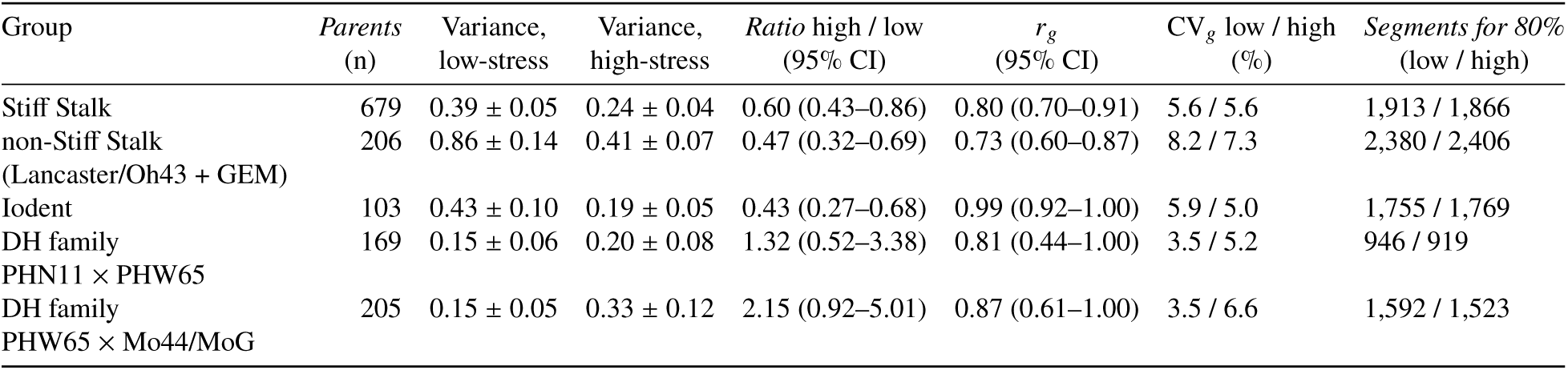
Testcross additive variance by group and condition, from the model with group deviations, in (t/ha)^2^ (± standard error). The *Ratio* column is the high-stress variance divided by the low-stress variance, with the 95% CI calculated on the log scale. *r_g_* is the genetic correlation between conditions, with the 95% CI truncated at 1. CV*_g_* is the genetic coefficient of variation, the genetic standard deviation divided by the mean of the hybrids of the model in that condition. The *Parents* column gives the number of parents of the group in the model, and the *Segments for 80%* column gives the number of 164-kb segments, out of 10,046, needed to account for 80% of the segment variance of the group.

| Group | *Parents*<br>(n) | Variance,<br>low-stress | Variance,<br>high-stress | *Ratio* high / low<br>(95% CI) | *r*<sub>*g*</sub><br>(95% CI) | CV<sub>*g*</sub> low / high<br>(%) | *Segments for 80%*<br>(low / high) |
|---|---|---|---|---|---|---|---|
| Stiff Stalk | 679 | 0.39 ± 0.05 | 0.24 ± 0.04 | 0.60 (0.43–0.86) | 0.80 (0.70–0.91) | 5.6 / 5.6 | 1,913 / 1,866 |
| non-Stiff Stalk<br>(Lancaster/Oh43 + GEM) | 206 | 0.86 ± 0.14 | 0.41 ± 0.07 | 0.47 (0.32–0.69) | 0.73 (0.60–0.87) | 8.2 / 7.3 | 2,380 / 2,406 |
| Iodent | 103 | 0.43 ± 0.10 | 0.19 ± 0.05 | 0.43 (0.27–0.68) | 0.99 (0.92–1.00) | 5.9 / 5.0 | 1,755 / 1,769 |
| DH family<br>PHN11 × PHW65 | 169 | 0.15 ± 0.06 | 0.20 ± 0.08 | 1.32 (0.52–3.38) | 0.81 (0.44–1.00) | 3.5 / 5.2 | 946 / 919 |
| DH family<br>PHW65 × Mo44/MoG | 205 | 0.15 ± 0.05 | 0.33 ± 0.12 | 2.15 (0.92–5.01) | 0.87 (0.61–1.00) | 3.5 / 6.6 | 1,592 / 1,523 |

The genetic correlation between conditions was below 1 in the non-Stiff Stalk group (0.73; 95% CI 0.60–0.87) and in the Stiff Stalk group (0.80; 0.70–0.91). In the Iodent group (0.99; 0.92–1.00) and in the two DH families, the interval includes 1 (Table 1).

The DH families have the smallest variance under low stress (0.15 and 0.15) and wide confidence intervals under high stress, where 71% and 73% of their observations are in only two environments.

### 3.4 Variance across the genome

The additive variance of each group is distributed across the genome (Figure 4). Under low stress, 1,913 of the 10,046 164-kb segments are needed to account for 80% of the segment variance in the Stiff Stalk group, 2,380 in the non-Stiff Stalk group and 1,755 in the Iodent group. Under high stress, these numbers are 1,866, 2,406 and 1,769 (Table 1). In the three heterotic groups, no segment reaches 1% of the segment variance of its group.

**Figure 4.**
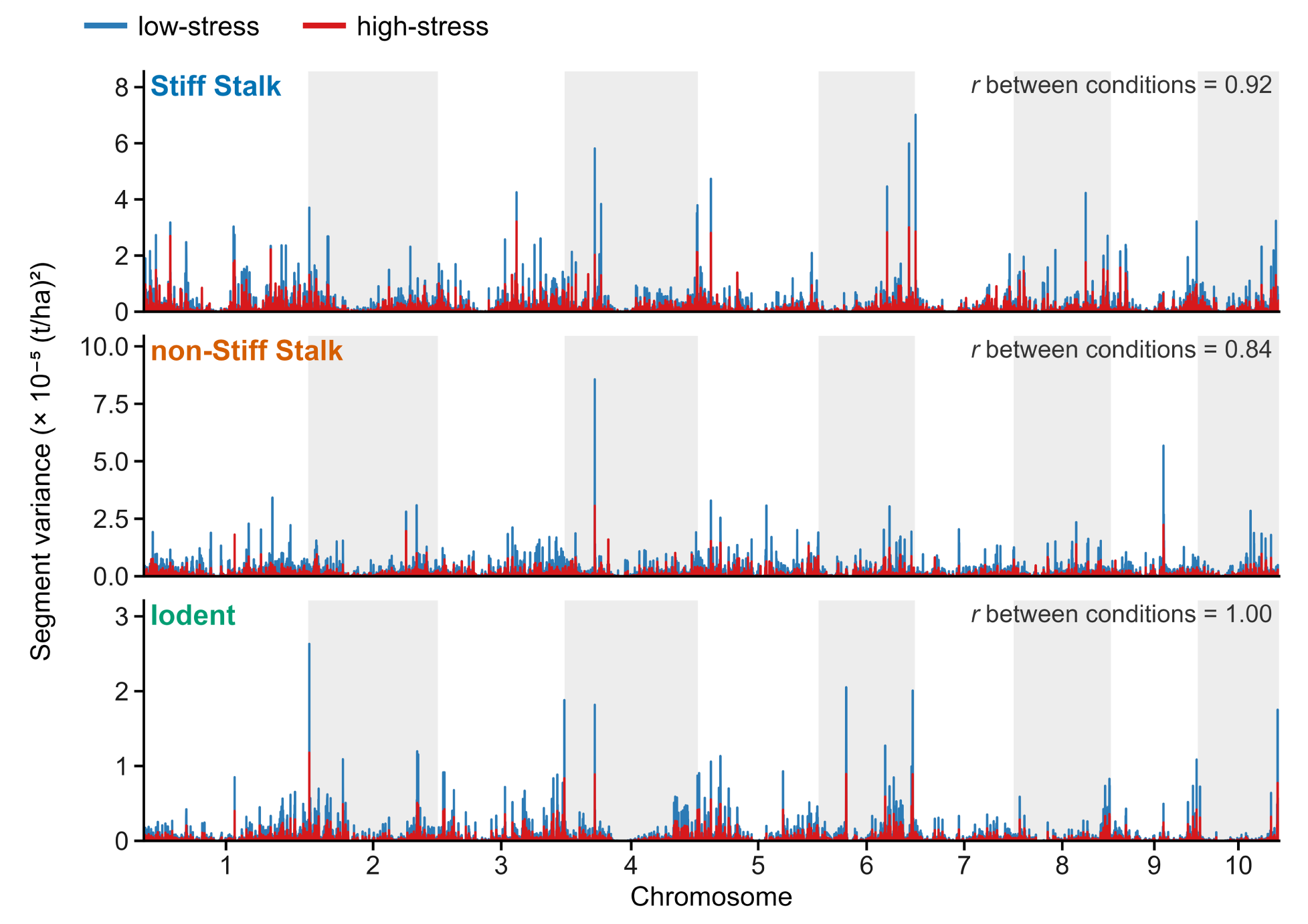
Additive variance across the genome in the three heterotic groups. Each panel is one group, with its own y-axis scale. The x-axis is the genome, in 10,046 164-kb segments, with the chromosomes in alternating bands. The y-axis is the segment variance, that is, the variance of the local GEBVs among the parents of the group in the model, under low stress (blue) and under high stress (red). *r* is the Pearson correlation between the segment variances of the two conditions. The two DH families are shown in Supplementary Figure S3.

The distribution of variance coincided only partly among the groups. The Spearman correlation between the segment variances of the three groups was 0.62–0.63 under low stress and 0.61–0.64 under high stress. Of the segments in the top 20% for variance in one group, 44–49% were also in the top 20% of another group (Figure 4).

Within each group, the distribution was similar in both conditions, with lower values under high stress. The Pearson correlation between the low- and high-stress segment variances was 0.92 in the Stiff Stalk group, 0.84 in the non-Stiff Stalk group and 1.00 in the Iodent group. Of the segments in the top 20% for variance, 83%, 74% and 97% coincided in the two conditions (Supplementary Table S5). The segments that moved up or down in the variance ranking between conditions were also not the same from one group to another, with a Spearman correlation of −0.01 to 0.03 between groups for the change in ranking.

There is G×E within groups, but it is spread over many segments, each with little variance. The correlation of local GEBVs between the two conditions (*r _j_*) had a median of 0.99 in the Stiff Stalk group, 0.96 in the non-Stiff Stalk group and 1.00 in the Iodent group. Segments with *r _j_* below 0.8 were 14% of the segments in the Stiff Stalk group, 23% in the non-Stiff Stalk group and 0.5% in the Iodent group. In the Stiff Stalk group, they account for 2.4% of the segment variance of the group under low stress and 2.1% under high stress. In the non-Stiff Stalk group, they account for 6.8% and 7.2%. Among the segments in the top 20% for variance in both conditions, only 0.8% in the Stiff Stalk group and 2.1% in the non-Stiff Stalk group had *r _j_* below 0.8, and none in the Iodent group (Supplementary Table S5). The segment variance of the two DH families is shown in Supplementary Figure S3.

### 3.5 Stacking ceiling

The ratio of the stacking ceiling to the best evaluated hybrid depends on segment resolution, not on condition (Figure 5a). At 164 kb, the ceiling was 31.6 t/ha under low stress and 23.3 t/ha under high stress, 117% and 105% above the best evaluated hybrid in each condition (14.6 and 11.4 t/ha). With the highest local GEBV of each segment in place of the 99th percentile, or without the two DH families in the non-Stiff Stalk pool, the ratio varied by at most 12 percentage points (Supplementary Table S2).

**Figure 5.**
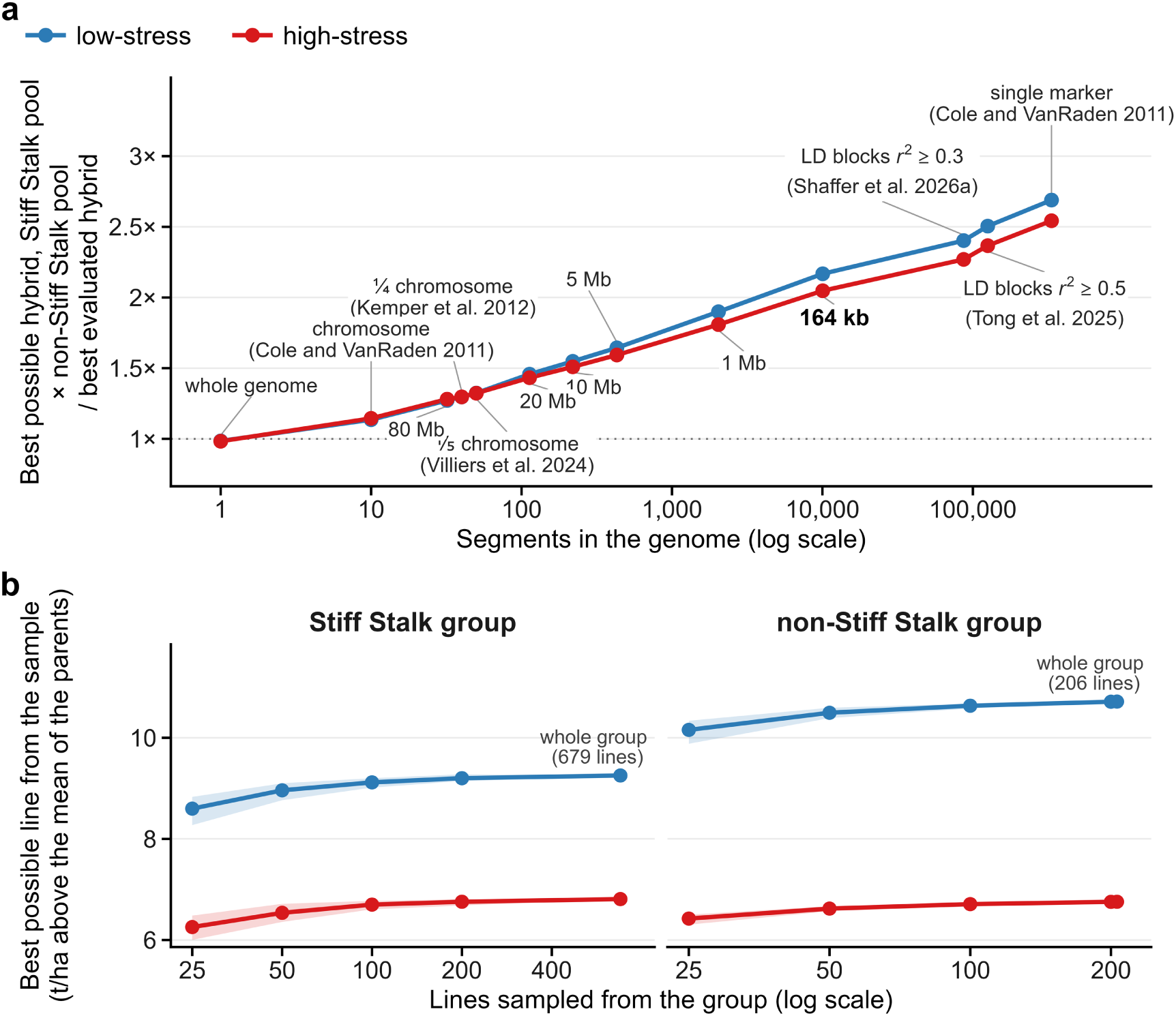
Stacking ceiling and segment resolution, under low stress (blue) and under high stress (red). Panel a shows the ratio of the ceiling, which is the cross between the best possible lines of the Stiff Stalk and non-Stiff Stalk pools, to the best evaluated hybrid, by the number of segments into which the genome is divided (log scale), from the whole genome to the single marker. All points were calculated with these data and the same marker effects, and the line only connects the points. The studies in parentheses indicate where each resolution was used, not their results. The best evaluated hybrid, among the hybrids of the model that cross the two pools, was PHW52 × NK787 under low stress (14.6 t/ha) and PHN11_PHW65_0294 × CG102 under high stress (11.4 t/ha). Panel b shows the best possible line in samples of *n* lines from the Stiff Stalk and non-Stiff Stalk groups, at 164 kb, in t/ha above the mean of the parents of the model. The line is the mean of 30 samples, the band is the range, and the last point is the whole group.

At the resolutions used in the literature, the ceiling was 14–15% above the best evaluated hybrid with whole chromosomes and 154–169% above with single markers, the lower and upper limits of Cole and VanRaden (2011). With one quarter or one fifth of a chromosome (Kemper et al. 2012; Villiers et al. 2024), it was 30–32% above, and with LD blocks (Shaffer et al. 2026a; Tong et al. 2025), 127–151% above.

The ceiling depended little on the number of lines. With samples of only 100 lines, the best possible line of the Stiff Stalk group and that of the non-Stiff Stalk group already reached 98–99% of the value obtained with the whole group (Figure 5b). With 100 phenotyped lines in each group, the best possible line of the non-Stiff Stalk group had, under low stress, a higher value (10.6 t/ha above the mean of the parents of the model) than that of the Stiff Stalk group (9.1) and that of the Iodent group (5.4). Under high stress, those of the non-Stiff Stalk and Stiff Stalk groups were equal (6.7 t/ha), and that of the Iodent group was lower (3.6).

### 3.6 Founder selection

With 25 founders, the stacking set reached 95% of the potential of the whole Stiff Stalk pool (9.27 of 9.80 t/ha above the mean of the parents of the model, under low stress) and 94% of that of the whole non-Stiff Stalk pool (10.84 of 11.51 t/ha), and, under high stress, 95% and 94%. Truncation reached 56–92% (Figure 6a; Supplementary Table S6). With 8 and 50 founders, stacking reached 78–82% and 97%. With 4 to 50 founders, the advantage of stacking over truncation was largest with 8 to 20 founders. It grew with the mean coancestry of the truncation founders: the potential of stacking was 1–4% higher than that of truncation when this coancestry was 0.05–0.09 and 36–82% higher when it was 0.32–0.47 (Supplementary Table S6). The results below are for 25 founders.

**Figure 6.**
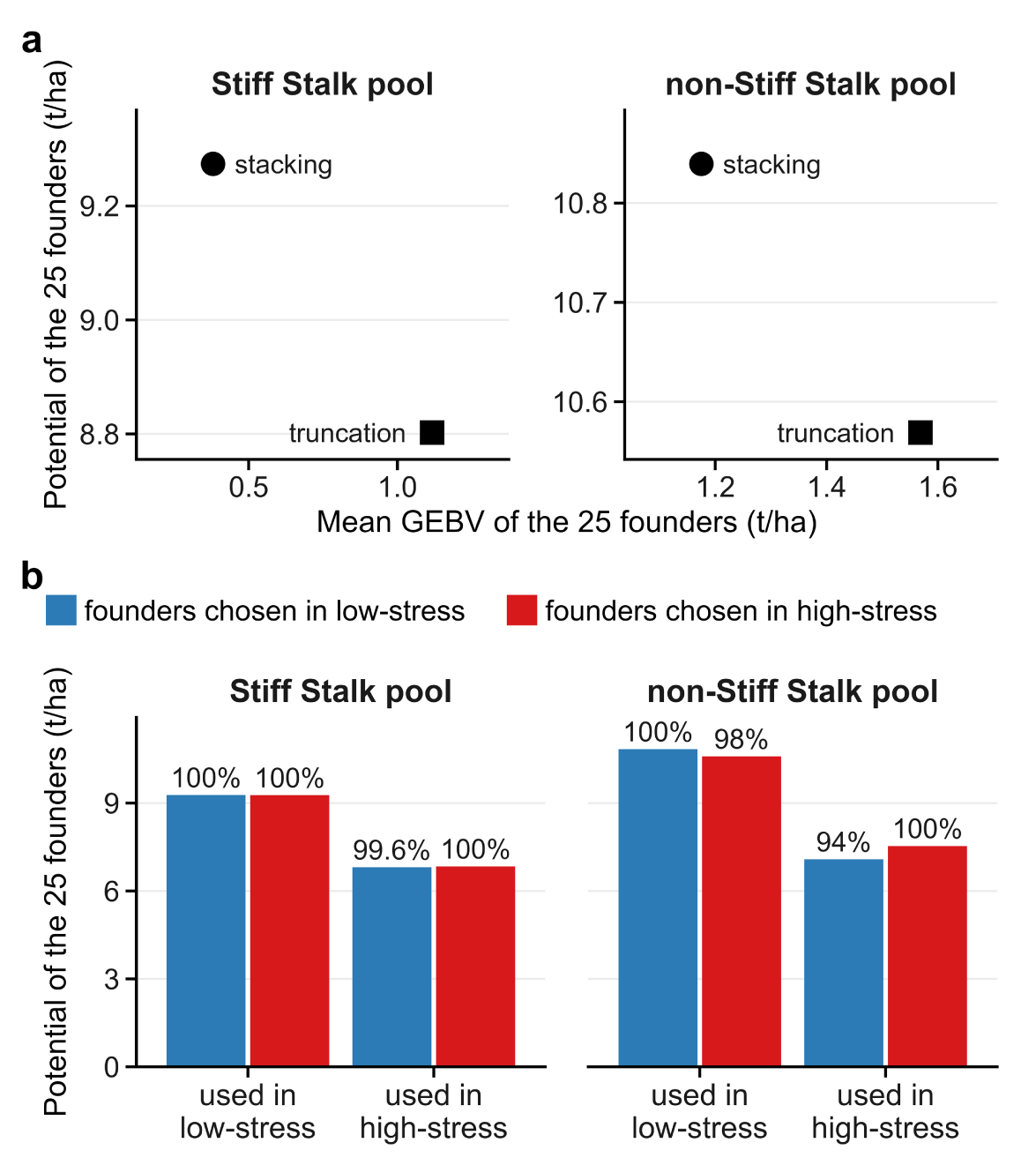
Selection of 25 founders in the Stiff Stalk and non-Stiff Stalk pools, in t/ha above the mean of the parents of the model. The potential (*Potential of the 25 founders*) is the sum, over the 10,046 164-kb segments, of the mean of the two highest local GEBVs from different founders in the set, and the mean GEBV (*Mean GEBV of the 25 founders*) is the mean of the GEBVs of the founders. Panel a compares stacking and truncation under low stress; the results for high stress are in Supplementary Table S6. Panel b shows the potential of each set in the two conditions and the retention, as a percentage of the potential of the set chosen in the same condition.

Stacking reached higher potential with founders of lower mean GEBV. Under low stress, the mean GEBV of its founders was 0.74 t/ha lower than that of the truncation founders in the Stiff Stalk pool and 0.39 t/ha lower in the non-Stiff Stalk pool. The stacking founders were more diverse. The mean coancestry of the sets was 0.03–0.06 for stacking, compared with 0.06–0.38 for truncation, while the mean over all pairs of lines in the pool is 0.15 in the Stiff Stalk pool and 0.09 in the non-Stiff Stalk pool. On average over the segments, the 25 stacking founders have 8–10 different versions per segment, compared with 3–9 for truncation. For truncation, the most common version of a segment appears in 14–15 of the 25 founders in the Stiff Stalk pool, compared with 11 for stacking (24–33% more). In the non-Stiff Stalk pool, it appears in 10–17, compared with 9–10 (9–67% more) (Supplementary Table S6).

Retention was 99.6–100% in the Stiff Stalk pool and 94–98% in the non-Stiff Stalk pool (Figure 6b), although the sets of the two conditions have fewer than half of their founders in common (12 of 25 in the Stiff Stalk pool and 7 of 25 in the non-Stiff Stalk pool).

### 3.7 Genome-wide association and segments

None of the loci associated with grain yield was detected in both conditions, either at the Bonferroni threshold or at a 5% FDR. At the Bonferroni threshold, the GWAS detected 25 loci under low stress and 18 under high stress (Figure 7a), and 16 in the delta analysis (Supplementary Figure S4; Supplementary Table S4). At a 5% FDR, there were 36 loci under low stress (37 SNPs), 21 under high stress (23 SNPs) and 19 in the delta analysis (19 SNPs; Supplementary Table S3).

**Figure 7.**
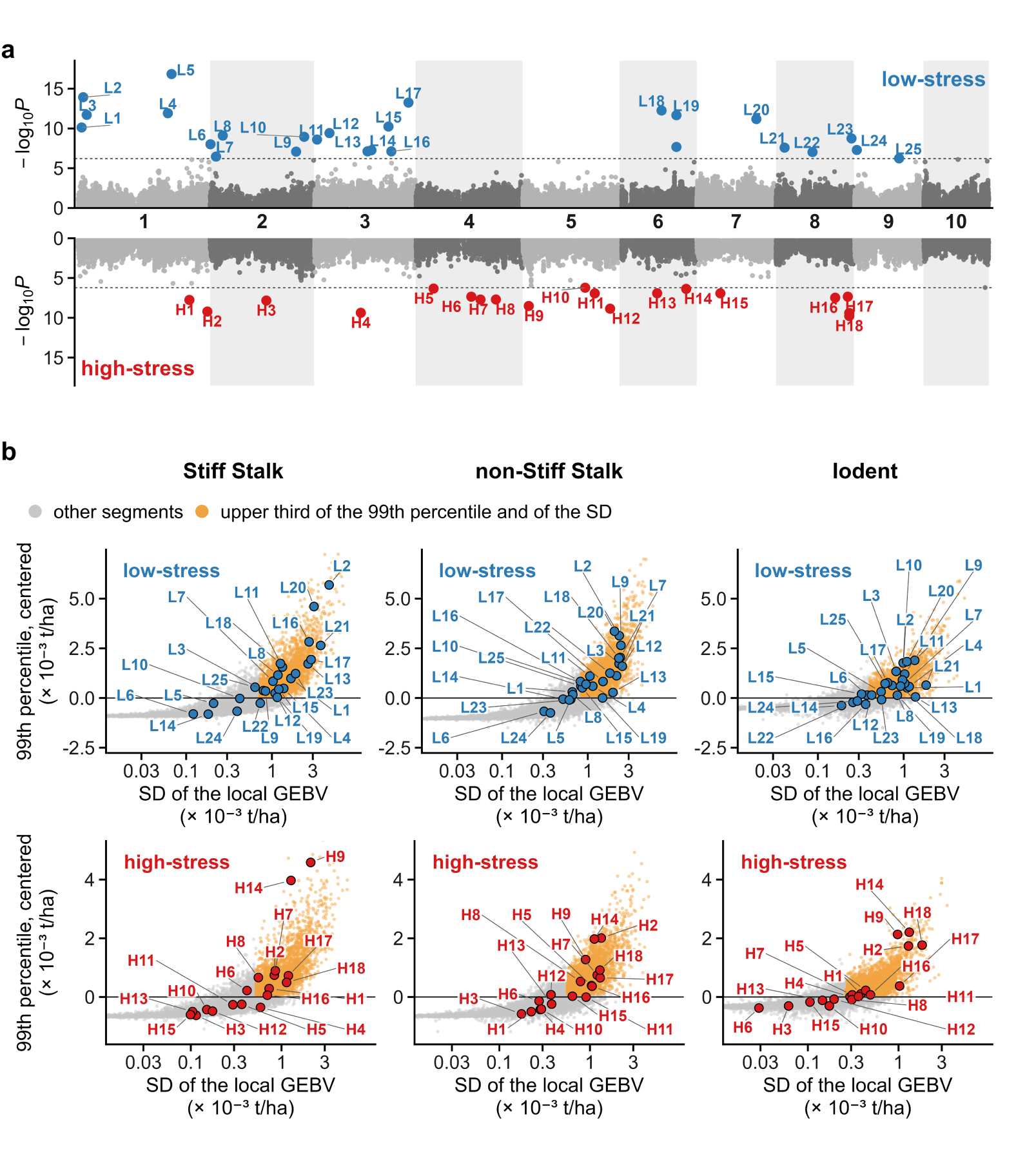
GWAS loci and 164-kb segments, under low stress (blue) and under high stress (red). Panel a shows the GWAS of grain yield under low stress (top) and under high stress (bottom). Each point is a SNP, with the lower *P*-value of BLINK and FarmCPU, and only 15% of the SNPs with *P* > 0.1 were plotted. The dashed line is the Bonferroni threshold for the number of independent tests of each condition (section 2.11). The colored SNPs, which reach it with at least one of the two methods, form the loci in Supplementary Table S4, numbered L1 to L25 under low stress and H1 to H18 under high stress. Panel b shows the segments of each heterotic group, in each condition. The x-axis is the standard deviation (SD) of the local GEBV among the parents of the group in the model, on a log scale. The y-axis is the 99th percentile of the local GEBV of the segment among the same parents, minus the mean of the 99th percentiles of all segments. In orange are the high-effect, high-variance segments (section 2.11), those in the upper third for the 99th percentile and, at the same time, in the upper third for the standard deviation of the group in the condition (about 27% of the segments). The upper third for the standard deviation is the same as for the variance. Each locus is placed in the segment of its lead SNP, with the number from panel a. The axes were cut to show the point cloud, and 1–6% of the segments of each panel fall outside.

Each low-stress and high-stress locus had a smaller effect in the other condition (Supplementary Table S4). The favorable allele of the low-stress loci had an effect of 0.23–0.71 t/ha under low stress and −0.07 to 0.30 t/ha under high stress, with the same sign in both conditions for 23 of the 25. The favorable allele of the high-stress loci had an effect of 0.25–0.55 t/ha under high stress and −0.16 to 0.30 t/ha under low stress.

None of the lead SNPs of the delta loci reached the threshold in the low- or high-stress GWAS. For 13 of the 16, the allele that increases the delta had a positive effect under low stress and a negative effect under high stress, with small effects in both conditions (Supplementary Table S4). Two delta loci lie less than 1 kb from the lead SNP of a low- or high-stress locus: D8 (chr3:180.34), near the low-stress locus L16, and D9 (chr4:129.83), near the high-stress locus H6 (Figure 7a; Supplementary Figure S4).

The loci were not concentrated in the segments that lose the least variance under stress. Among the 1,362 parents of the model, the ratio of the standard deviation of the local GEBV under high stress to that under low stress had a median of 0.75 across the 164-kb segments. Of the 18 high-stress loci, 7 are in the third with the highest ratio (above 0.83), 7 in the middle third and 4 in the third with the lowest ratio (below 0.68). The 25 low-stress loci were split evenly among the three thirds (8, 9 and 8) (Supplementary Table S4).

Many loci lie in the high-effect, high-variance segments (Figure 7b). These segments are about 27% of the segments of each group, in each condition. Under low stress, they contain 17 of the 25 loci in the Stiff Stalk group, 15 in the non-Stiff Stalk group and 18 in the Iodent group. Under high stress, they contain 9 of the 18 loci in each of the three groups (Supplementary Table S4). In the Stiff Stalk, non-Stiff Stalk and Iodent groups, 84%, 78% and 95% of these segments coincided in the two conditions.

## Discussion

High stress reduced testcross additive variance by 40–57% in the three heterotic groups. Within each group, the distribution of variance across the genome was similar in both conditions, with lower values under high stress. The ranking of parents, however, changed partly in the Stiff Stalk and non-Stiff Stalk groups. The GWAS loci were specific to each condition. They lay, to a large extent, in the high-effect, high-variance segments. Unlike the loci, these segments were nearly the same in both conditions. In stacking, the ratio of the ceiling to the best evaluated hybrid was nearly the same in both conditions. The set of founders chosen in one condition retained, in the other, almost all of the potential of the set chosen there.

### Variance by group and between conditions

Under low stress, the non-Stiff Stalk group had about twice the additive variance of the Stiff Stalk group. In G2F single-cross hybrids, analyzed without separating conditions, the general combining ability variance of the non-Stiff Stalk group for grain yield was about twice that of the Stiff Stalk group (Godoy dos Santos et al. 2026). Under high stress, variance decreased in all three groups, as did the variance among hybrids under managed drought (Cooper et al. 2014) and the genetic variability in the lower-yielding G2F environments (Tolley et al. 2023).

In the Stiff Stalk group, variance decreased by 40% and the genetic coefficient of variation did not change (5.6% in both conditions), that is, the decrease followed that of grain yield. In the non-Stiff Stalk and Iodent groups, variance decreased by 53% and 57%. The genetic coefficient of variation also decreased, from 8.2% to 7.3% and from 5.9% to 5.0%, that is, variance decreased more than the decrease in grain yield would explain. The percentage of decrease, however, did not differ significantly among the groups (*P* ≥ 0.24), and the non-Stiff Stalk group still had the largest variance under high stress. The ranking of parents also partly changed between conditions, with a genetic correlation below 1 in the non-Stiff Stalk group (0.73) and in the Stiff Stalk group (0.80), and this re-ranking affects the choice of parents (Cooper and Messina 2023).

### Segments and loci

Additive variance is distributed across the genome, and 1,755–2,406 segments of 164 kb, depending on the group and condition, are needed to account for 80% of the segment variance. The highest-variance segments of one group coincided only partly with those of another, and the segments that moved up or down in the variance ranking between conditions were not the same from one group to another. In Dent × Flint maize, marker effects also differed between heterotic groups (Technow et al. 2014). Within each group, the distribution was similar in both conditions, as in wheat, in which high-variance blocks for grain yield were observed under different levels of nitrogen fertilization and fungicide use (Voss-Fels et al. 2019). This occurred even with the grid chosen to maximize the contrast between conditions, a contrast that may be smaller in new environments. In most segments, the ranking of parents was nearly the same in both conditions, and G×E appeared mainly in low-variance segments. The decrease in variance under stress, however, was not the same in all segments. In some segments it decreased more, and other segments kept their variance.

No locus associated with grain yield was common to both conditions, and each locus had a smaller effect in the condition in which it was not identified. This means that no locus passed the threshold in both conditions, not that common loci do not exist. GWAS detects the larger effects, and power differs between conditions: there are 2,756 hybrids under low stress, where the variance is larger, and 2,554 under high stress. The loci are also candidates. The union of BLINK and FarmCPU detects the loci but does not confirm them; confirmation requires independent data. The GWAS of the delta, which tests the difference between conditions directly, found loci with small effects, and for most of them the same allele increased grain yield under low stress and reduced it under high stress. In maize testcross hybrids in Europe, almost all grain yield loci also had scenario-dependent effects. Tolerance loci had a strong effect only in the hot or dry scenarios, and sensitivity loci lost their effect under stress (Millet et al. 2016). In tropical maize testcross hybrids, several associations were detected only under water stress (Anoni et al. 2026). In G2F, in 220 DH lines crossed with three testers, only 2 of the 96 grain yield QTL were shared across environments or testers (Ozair et al. 2025). Thus, a similar distribution of variance does not mean that the same genetic basis acts in both conditions. A 164-kb segment can have high variance in both for different reasons, such as different variants within it or the same variant with a different effect.

The loci found marker by marker lay, to a large extent, in the high-effect, high-variance segments of each group, more under low stress (15–18 of the 25 loci, depending on the group) than under high stress (9 of the 18). Thus, GWAS and local GEBVs pointed to the same regions, although the former tests one marker at a time and the latter spreads the effect over the segment. The loci were not concentrated in the segments that lose the least variance under stress. In wheat and barley, the highest-variance blocks also coincided with known QTL, for stripe rust resistance and for spike row type (Tong et al. 2025; Shaffer et al. 2026b).

### Stacking ceiling

The stacking ceiling is a selection limit and corresponds to the predicted value of the cross between the best possible lines of the two pools, considering only additive values. It is a theoretical limit: it assumes that the best segments can be combined by recombination, without linkage drag or epistasis, and it does not represent the gain expected in a few cycles. In the same panel and with the same marker effects, the ratio of the ceiling to the best evaluated hybrid depended on segment resolution. It was nearly the same in both conditions, because the ceiling and the best evaluated hybrid decreased in similar proportions under high stress. The ceiling was 14–15% above the best evaluated hybrid with whole chromosomes and 154–169% above with single markers, which correspond to the lower and upper limits of Cole and VanRaden (2011). Published ceilings range from 1.23 times the best available line, for grain yield in wheat, to 6 times the best sire, for dairy economic value, and reflect the diversity of each population (Hayes et al. 2024). These values were also obtained at different scales and resolutions. In a biparental maize cross, with only two parents, the best possible line with one or two targeted recombinations per chromosome had 7–10% higher testcross performance than the best of the 180 lines from the cross (Bernardo 2017). With the 679 Stiff Stalk pool parents and the 683 non-Stiff Stalk pool parents of this study, the ceiling already exceeded the best evaluated hybrid with whole chromosomes, without recombination within them.

### Founders

With 25 founders, stacking, which chooses the set with the highest potential, reached 94–95% of the potential of all parents in the pool, more than truncation, even with founders whose mean GEBV was 0.39–0.74 t/ha lower under low stress. This is because stacking chooses parents that complement each other, and a parent can enter the set without being among the best by total GEBV, provided it has high local GEBVs in some segments (Kemper et al. 2012). The advantage of stacking is larger when the best lines by total GEBV are related, because related lines carry the same versions of many segments. As in Villiers et al. (2024), stacking resulted in a more diverse base, with lower coancestry and more versions of each segment. In wheat, the mean of the truncation founders was also about 0.30 t/ha higher than that of the stacking founders (Shaffer et al. 2026a).

Between conditions, the sets had different compositions and similar potential. The sets of the two conditions had 12 of 25 founders in common in the Stiff Stalk pool and 7 of 25 in the non-Stiff Stalk pool. Even so, the set chosen in one condition, when used in the other, had at most 0.4% less potential than the set chosen there in the Stiff Stalk pool, and 2–6% less in the non-Stiff Stalk pool.

### Use in breeding programs

Historically, high stress has been handled differently across regions of the Corn Belt. In the western region, which is drier and has more irrigation (Ting et al. 2023), direct selection for grain yield under drought began in the 1950s, whereas in the central region the gain under drought resulted from the evaluation of hybrids in many environments (Cooper et al. 2014). With the hotter and drier summers projected for the whole Corn Belt (Ting et al. 2021), high stress should become more important for programs in the central and eastern regions as well. In the G2F network, environmental components related to water stress have already been shown to be important for enviromic prediction (Godoy Filho et al. 2026a).

In addition to predicting performance under stress, programs need to choose parents, donors and founders for these conditions. Most programs are ongoing and will not be restarted. In each cycle, they choose parents and can bring in new lines from outside the program as donors of segments with high local GEBVs (Allier et al. 2020). For these programs, the choice of donor lines can consider the high-effect, high-variance segments of each group, which were nearly the same in both conditions and where many loci lay. The loci of each condition can help identify which of these segments may be important for low- and high-stress years and regions.

New breeding programs can use stacking once, to choose the founders, and then follow the usual selection (Villiers et al. 2024), evaluating the incorporation of new material throughout the process. The sets chosen in the two conditions had similar potential but fewer than half of their founders in common, and the program can combine founders from both conditions to broaden the genetic variability of the base. In simulations, stacking maintained more diversity and had greater long-term genetic gain than truncation (Villiers et al. 2024; Shaffer et al. 2026a). With enough data to estimate QTL reliably, the GWAS loci could also be included in the stacking algorithm as additional information about the segments that contain them, an approach that has yet to be developed.

### Conclusion

In G2F, high stress reduced testcross additive variance in the three heterotic groups, and the distribution of this variance across the genome remained similar. The ranking of parents changed partly in the Stiff Stalk and non-Stiff Stalk groups, and the GWAS loci were specific to each condition. The loci lay, to a large extent, in the high-effect, high-variance segments. Unlike the loci, these segments were nearly the same in both conditions. The local GEBVs of these segments can guide the choice of donors, and the loci of each condition can help identify segments that may be important. The founders chosen in each condition have different compositions, and each set retained, in the other condition, almost all of the potential of the set chosen there. Combining founders from both conditions broadens the genetic variability of the base. Choosing founders by stacking also resulted in a more diverse base, which tends to favor greater long-term gains than truncation on total GEBV. With the hotter and drier summers projected for the Corn Belt, these choices should become more important.

## Data availability

All data needed to reproduce the results of this study, including phenotypic, genomic and enviromic data, are available on Zenodo under DOI https://doi.org/10.5281/zenodo.23689239. The data are provided as .csv and .rds files, and the repository also includes the R code that reproduces the analyses.

## Acknowledgments

We gratefully acknowledge the Department of Genetics, Luiz de Queiroz College of Agriculture, University of São Paulo (ESALQ/USP), for its institutional support. During the preparation of this work, the authors used large language models (Google Gemini and Anthropic Claude) for coding support/debugging and text refinement.

## Supplementary material

Supplementary Figures S1–S4 and Tables S1–S6 are in the Supplementary Material file, and Supplementary File S1 is in File_S1.xlsx.

