## Supplementary Material. for "Local GEBVs and additive variance in maize heterotic groups across low- and high-stress environments"

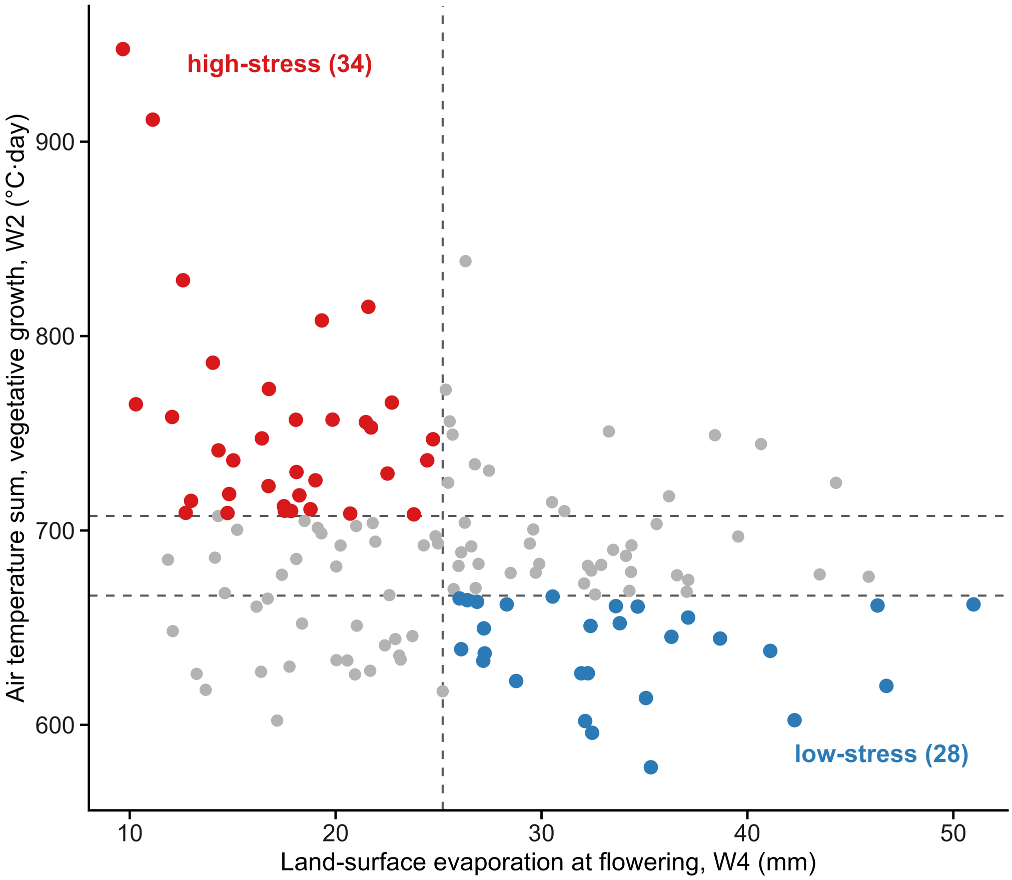


**Supplementary Figure S1.** Climate grid of the rainfed environments, with land-surface evaporation at flowering (W4) on the x-axis and air temperature sum during vegetative growth (W2) on the y-axis. The vertical line is the evaporation median, and the horizontal lines are the temperature terciles. Blue points are the low-stress environments (high evaporation and coldest tercile), red points are the high-stress environments (low evaporation and warmest tercile), and gray points are the environments in the other cells, which are not used in the analyses by condition.

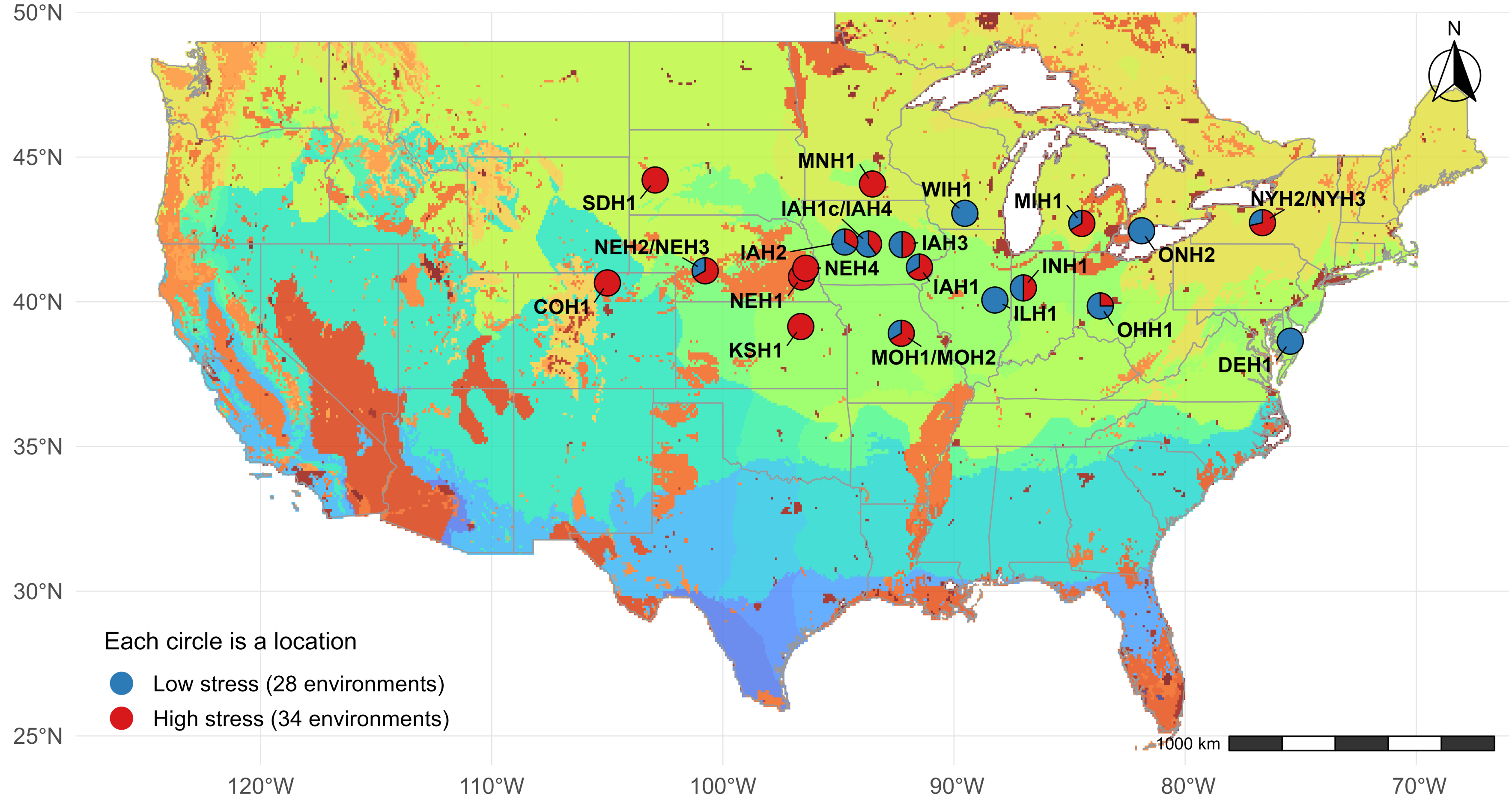


**Supplementary Figure S2.** Locations of the 62 environments over the FAO agro-ecological zones (GAEZ; FAO 2025). Each circle is a location, split according to the number of low-stress (blue, 28 in total) and high-stress (red, 34 in total) environments evaluated there. Locations less than 0.3° apart are shown as a single circle, with their names separated by a slash.

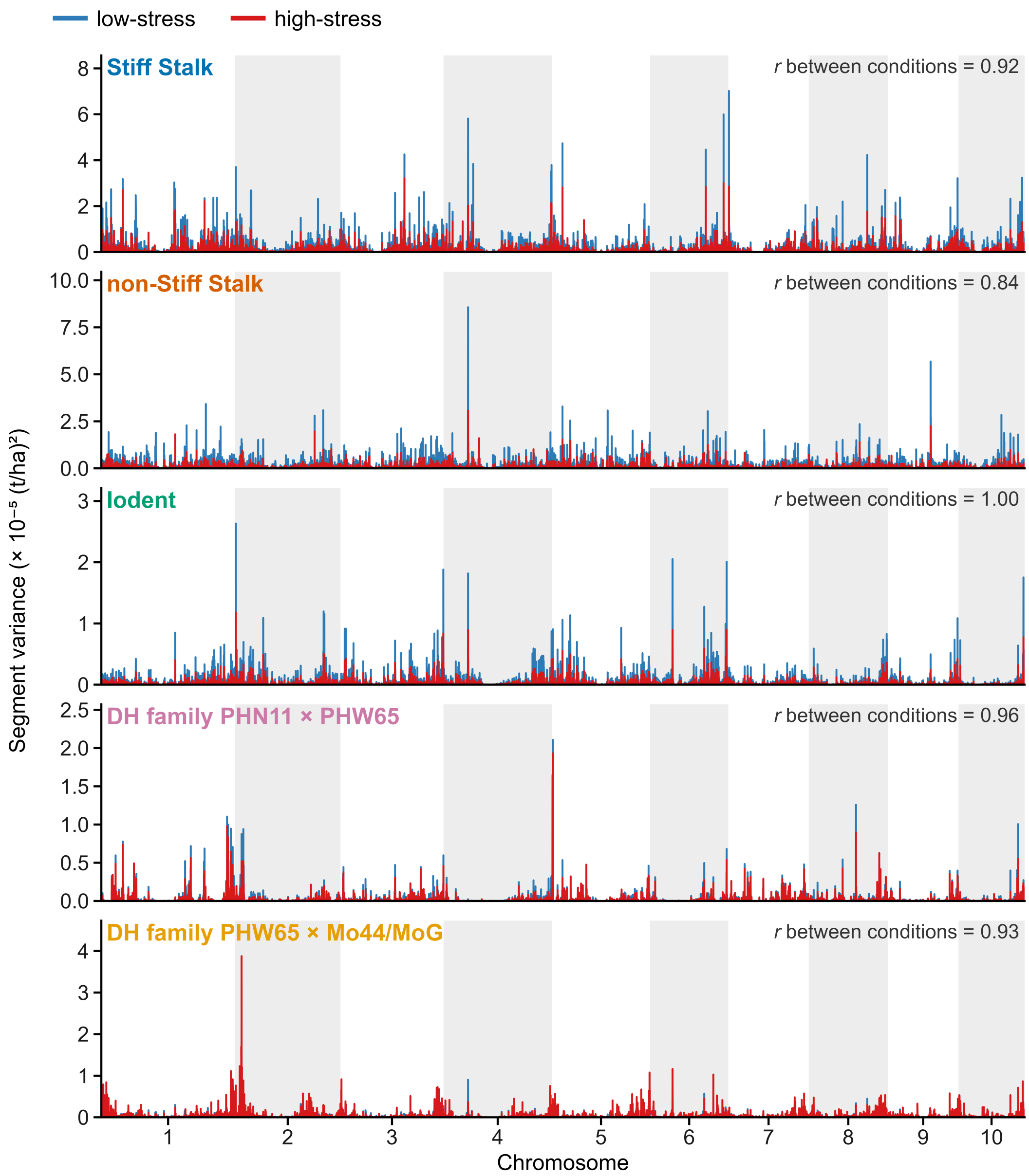


**Supplementary Figure S3.** Additive variance across the genome in the three heterotic groups and the two DH families, as in Figure 4. In the families, stretches without variance are segments in which the two parents of the family are identical.

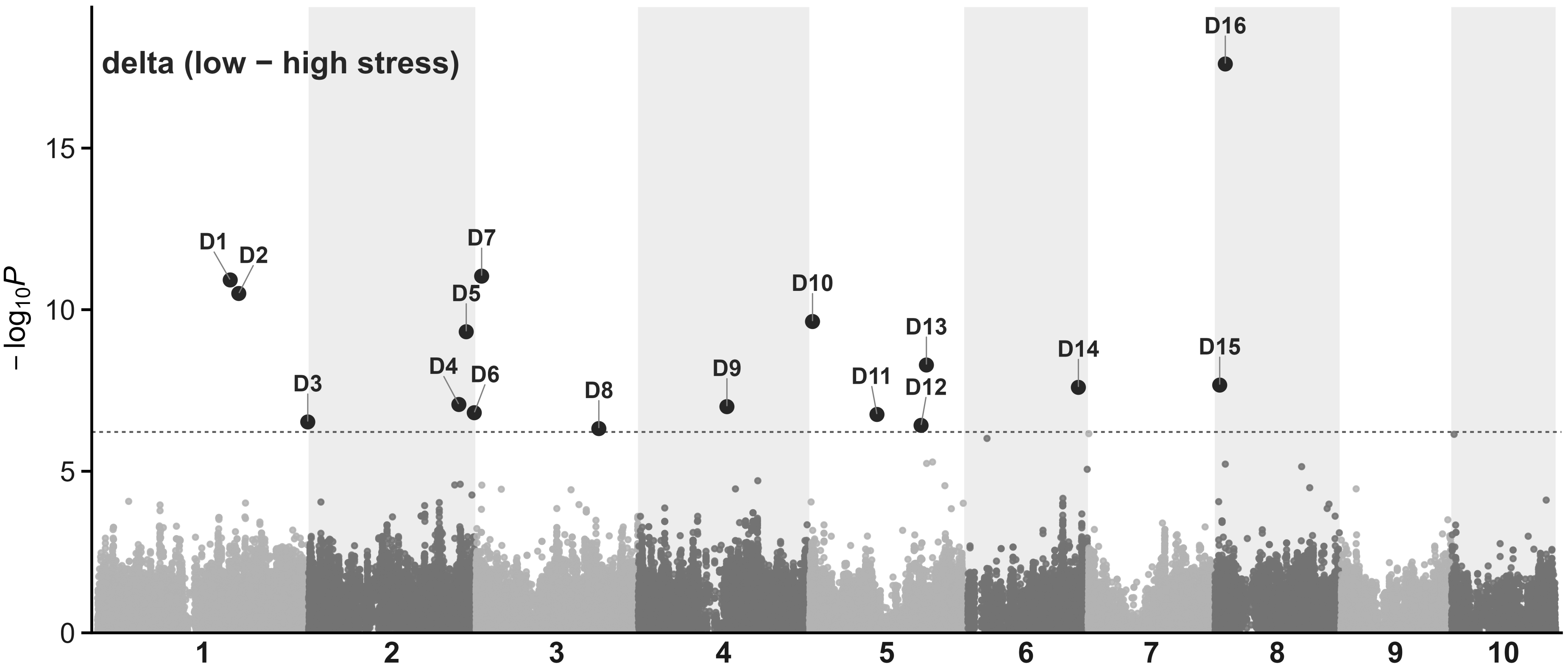


**Supplementary Figure S4.** GWAS of the delta (low − high stress), in the 2,084 hybrids evaluated in both conditions. Each point is a SNP, with the lower $P$-value of BLINK and FarmCPU, and only 15% of the SNPs with $P > 0.1$ were plotted. The dashed line is the Bonferroni threshold for the number of independent tests of the delta (section 2.11). The highlighted SNPs, which reach it with at least one of the two methods, form the delta loci in Supplementary Table S4, numbered D1 to D16.

**Supplementary Table S1:** The 62 environments, with condition, year, coordinates, the two grid covariates (window total) and the number of hybrids evaluated.

| Environment | Condition | Year | Latitude | Longitude | Land-surface evaporation at flowering, W4 (mm) | Air temperature sum, vegetative growth, W2 (°C·day) | Hybrids evaluated (n) |
|---|---|---|---|---|---|---|---|
| DEH1_2016 | low-stress | 2016 | 38.64 | −75.45 | 34.7 | 661 | 240 |
| IAH1_2019 | low-stress | 2019 | 41.20 | −91.50 | 30.5 | 666 | 596 |
| IAH1c_2014 | low-stress | 2014 | 42.00 | −93.70 | 33.8 | 652 | 137 |
| IAH2_2014 | low-stress | 2014 | 42.07 | −94.72 | 37.1 | 655 | 239 |
| IAH2_2019 | low-stress | 2019 | 42.07 | −94.72 | 38.7 | 644 | 597 |
| IAH3_2015 | low-stress | 2015 | 41.98 | −92.24 | 33.6 | 661 | 426 |
| IAH3_2017 | low-stress | 2017 | 41.98 | −92.24 | 31.9 | 627 | 329 |
| IAH3_2019 | low-stress | 2019 | 41.98 | −92.24 | 41.1 | 638 | 591 |
| IAH4_2015 | low-stress | 2015 | 42.00 | −93.70 | 46.8 | 620 | 426 |
| IAH4_2019 | low-stress | 2019 | 42.00 | −93.70 | 27.2 | 633 | 1187 |
| ILH1_2018 | low-stress | 2018 | 40.06 | −88.23 | 51.0 | 662 | 400 |
| INH1_2017 | low-stress | 2017 | 40.48 | −87.00 | 32.1 | 602 | 178 |
| MIH1_2020 | low-stress | 2020 | 42.71 | −84.47 | 28.8 | 623 | 416 |
| MOH2_2014 | low-stress | 2014 | 38.93 | −92.35 | 42.3 | 602 | 247 |
| NEH2_2018 | low-stress | 2018 | 41.05 | −100.75 | 46.3 | 661 | 318 |
| NEH2_2019 | low-stress | 2019 | 41.05 | −100.75 | 26.1 | 639 | 356 |
| NYH2_2017 | low-stress | 2017 | 42.73 | −76.65 | 27.2 | 637 | 200 |
| NYH2_2018 | low-stress | 2018 | 42.73 | −76.65 | 26.0 | 665 | 401 |
| OHH1_2015 | low-stress | 2015 | 39.86 | −83.67 | 32.4 | 651 | 426 |
| OHH1_2018 | low-stress | 2018 | 39.86 | −83.67 | 35.1 | 614 | 389 |
| OHH1_2020 | low-stress | 2020 | 39.86 | −83.67 | 27.2 | 650 | 399 |
| ONH2_2015 | low-stress | 2015 | 42.45 | −81.88 | 26.4 | 664 | 357 |
| WIH1_2014 | low-stress | 2014 | 43.06 | −89.53 | 32.2 | 627 | 208 |
| WIH1_2016 | low-stress | 2016 | 43.06 | −89.53 | 26.9 | 663 | 320 |
| WIH1_2017 | low-stress | 2017 | 43.06 | −89.53 | 32.5 | 596 | 291 |
| WIH1_2018 | low-stress | 2018 | 43.06 | −89.53 | 36.3 | 645 | 412 |
| WIH1_2019 | low-stress | 2019 | 43.06 | −89.53 | 35.3 | 578 | 603 |
| WIH1_2020 | low-stress | 2020 | 43.06 | −89.53 | 28.3 | 662 | 396 |
| COH1_2017 | high-stress | 2017 | 40.65 | −105.00 | 10.3 | 765 | 222 |
| IAH1_2022 | high-stress | 2022 | 41.20 | −91.50 | 21.5 | 756 | 516 |
| IAH1_2023 | high-stress | 2023 | 41.20 | −91.50 | 17.8 | 710 | 514 |
| IAH2_2023 | high-stress | 2023 | 42.07 | −94.72 | 21.7 | 753 | 514 |
| IAH3_2016 | high-stress | 2016 | 41.98 | −92.24 | 23.8 | 708 | 392 |
| IAH3_2022 | high-stress | 2022 | 41.98 | −92.24 | 24.7 | 747 | 513 |
| IAH3_2023 | high-stress | 2023 | 41.98 | −92.24 | 16.7 | 723 | 510 |
| IAH4_2014 | high-stress | 2014 | 42.00 | −93.70 | 22.7 | 766 | 245 |
| IAH4_2022 | high-stress | 2022 | 42.00 | −93.70 | 12.1 | 758 | 510 |
| INH1_2014 | high-stress | 2014 | 40.48 | −87.00 | 9.7 | 948 | 242 |
| KSH1_2018 | high-stress | 2018 | 39.14 | −96.63 | 24.4 | 736 | 292 |
| MIH1_2022 | high-stress | 2022 | 42.71 | −84.47 | 17.5 | 710 | 511 |
| MIH1_2023 | high-stress | 2023 | 42.71 | −84.47 | 16.4 | 747 | 507 |
| MNH1_2014 | high-stress | 2014 | 44.07 | −93.53 | 18.2 | 718 | 236 |
| MNH1_2021 | high-stress | 2021 | 44.07 | −93.53 | 14.0 | 786 | 417 |
| MNH1_2022 | high-stress | 2022 | 44.07 | −93.53 | 14.3 | 741 | 435 |
| MNH1_2023 | high-stress | 2023 | 44.07 | −93.53 | 18.1 | 730 | 454 |
| MOH1_2023 | high-stress | 2023 | 38.90 | −92.21 | 16.8 | 773 | 520 |
| MOH2_2022 | high-stress | 2022 | 38.93 | −92.35 | 18.1 | 757 | 443 |
| NEH1_2020 | high-stress | 2020 | 40.86 | −96.60 | 22.5 | 729 | 248 |
| NEH1_2022 | high-stress | 2022 | 40.86 | −96.60 | 20.7 | 709 | 334 |
| NEH1_2023 | high-stress | 2023 | 40.86 | −96.60 | 21.6 | 815 | 333 |
| NEH2_2015 | high-stress | 2015 | 41.05 | −100.75 | 19.3 | 808 | 413 |

*Supplementary Table S1, continued*

| Environment | Condition | Year | Latitude | Longitude | Land-surface evaporation at flowering, W4 (mm) | Air temperature sum, vegetative growth, W2 (°C·day) | Hybrids evaluated (n) |
|---|---|---|---|---|---|---|---|
| NEH2_2021 | high-stress | 2021 | 41.05 | −100.75 | 12.6 | 829 | 248 |
| NEH2_2022 | high-stress | 2022 | 41.05 | −100.75 | 11.1 | 911 | 334 |
| NEH3_2015 | high-stress | 2015 | 41.09 | −100.78 | 14.8 | 719 | 412 |
| NEH4_2016 | high-stress | 2016 | 41.16 | −96.41 | 19.0 | 726 | 230 |
| NYH2_2015 | high-stress | 2015 | 42.73 | −76.65 | 19.9 | 757 | 356 |
| NYH2_2022 | high-stress | 2022 | 42.73 | −76.65 | 12.7 | 709 | 429 |
| NYH3_2015 | high-stress | 2015 | 42.73 | −76.65 | 15.0 | 736 | 358 |
| NYH3_2018 | high-stress | 2018 | 42.73 | −76.65 | 18.8 | 711 | 419 |
| NYH3_2022 | high-stress | 2022 | 42.73 | −76.65 | 13.0 | 715 | 436 |
| OHH1_2016 | high-stress | 2016 | 39.86 | −83.67 | 17.5 | 712 | 241 |
| SDH1_2015 | high-stress | 2015 | 44.21 | −102.93 | 14.8 | 709 | 322 |

**Supplementary Table S2:** Sensitivity of the stacking ceiling. The table gives the ratio of the ceiling to the best evaluated hybrid for each resolution and condition, with the 99th percentile of each segment, as in the text, with the highest local GEBV of each segment in place of the 99th percentile, and without the two DH families in the non-Stiff Stalk pool (309 lines of the non-Stiff Stalk and Iodent groups).

| Resolution | Segments (n) | Low-stress: main (p99) | Low-stress: maximum | Low-stress: without DH families | High-stress: main (p99) | High-stress: maximum | High-stress: without DH families |
|---|---|---|---|---|---|---|---|
| Whole genome | 1 | 0.99 | 1.07 | 1.00 | 0.98 | 1.06 | 0.93 |
| Chromosome | 10 | 1.14 | 1.24 | 1.16 | 1.15 | 1.23 | 1.07 |
| 80 Mb | 32 | 1.27 | 1.38 | 1.29 | 1.28 | 1.35 | 1.20 |
| 1/4 chromosome | 40 | 1.30 | 1.40 | 1.32 | 1.30 | 1.38 | 1.22 |
| 1/5 chromosome | 50 | 1.32 | 1.43 | 1.34 | 1.32 | 1.40 | 1.24 |
| 20 Mb | 113 | 1.46 | 1.56 | 1.48 | 1.43 | 1.50 | 1.36 |
| 10 Mb | 219 | 1.55 | 1.65 | 1.57 | 1.51 | 1.59 | 1.45 |
| 5 Mb | 430 | 1.64 | 1.76 | 1.66 | 1.59 | 1.68 | 1.53 |
| 1 Mb | 2037 | 1.90 | 2.01 | 1.92 | 1.81 | 1.90 | 1.75 |
| 164 kb | 10046 | 2.17 | 2.26 | 2.18 | 2.05 | 2.12 | 1.99 |
| LD blocks ($r^2 \geq 0.3$) | 86884 | 2.40 | 2.46 | 2.40 | 2.27 | 2.32 | 2.18 |
| LD blocks ($r^2 \geq 0.5$) | 125563 | 2.51 | 2.55 | 2.49 | 2.37 | 2.41 | 2.27 |
| Marker | 333382 | 2.69 | 2.72 | 2.67 | 2.54 | 2.57 | 2.42 |

**Supplementary Table S3:** SNPs significant at a false discovery rate (FDR) of 5% for each method and analysis, with position, effect (t/ha), *P*, FDR, whether the SNP reached the Bonferroni threshold (section 2.11) and the locus in Supplementary Table S4.

| Analysis | Method | SNP | Chr | Position (bp) | Effect (t/ha) | *P* | FDR (BH) | Bonferroni threshold | Locus (Table S4) |
|---|---|---|---|---|---|---|---|---|---|
| low-stress | BLINK | S1_7923994 | 1 | 7923994 | −0.474 | $7.57 \times 10^{-11}$ | $2.35 \times 10^{-6}$ | yes | chr1:7.92 |
| low-stress | BLINK | S1_11490086 | 1 | 11490086 | 0.448 | $1.19 \times 10^{-14}$ | $1.84 \times 10^{-9}$ | yes | chr1:11.49 |
| low-stress | BLINK | S1_19871213 | 1 | 19871213 | 0.467 | $1.99 \times 10^{-12}$ | $9.68 \times 10^{-8}$ | yes | chr1:19.87 |
| low-stress | BLINK | S1_209407708 | 1 | 209407708 | −0.559 | $1.22 \times 10^{-12}$ | $7.58 \times 10^{-8}$ | yes | chr1:209.41 |
| low-stress | BLINK | S1_218122062 | 1 | 218122062 | 0.646 | $1.46 \times 10^{-17}$ | $4.53 \times 10^{-12}$ | yes | chr1:218.12 |
| low-stress | FarmCPU | S1_218122062 | 1 | 218122062 | 0.354 | $1.34 \times 10^{-6}$ | 0.0321 | no | chr1:218.12 |
| low-stress | BLINK | S1_265325201 | 1 | 265325201 | −0.490 | $8.36 \times 10^{-7}$ | 0.0113 | no | |
| low-stress | BLINK | S1_288427007 | 1 | 288427007 | −0.190 | $1.04 \times 10^{-6}$ | 0.0134 | no | |
| low-stress | FarmCPU | S1_305384675 | 1 | 305384675 | −0.213 | $1.72 \times 10^{-6}$ | 0.0358 | no | |
| low-stress | BLINK | S2_880810 | 2 | 880810 | 0.361 | $9.68 \times 10^{-9}$ | 0.000215 | yes | chr2:0.88 |
| low-stress | BLINK | S2_13823291 | 2 | 13823291 | −0.229 | $3.46 \times 10^{-7}$ | 0.00513 | yes | chr2:13.82 |
| low-stress | BLINK | S2_29661916 | 2 | 29661916 | −0.280 | $3.43 \times 10^{-7}$ | 0.00513 | yes | chr2:29.66 |
| low-stress | FarmCPU | S2_29661916 | 2 | 29661916 | −0.386 | $7.87 \times 10^{-10}$ | 0.000176 | yes | chr2:29.66 |
| low-stress | BLINK | S2_201066037 | 2 | 201066037 | −0.244 | $8.24 \times 10^{-8}$ | 0.00142 | yes | chr2:201.07 |
| low-stress | BLINK | S2_219944439 | 2 | 219944439 | 0.562 | $1.15 \times 10^{-8}$ | 0.000238 | yes | chr2:219.94 |
| low-stress | FarmCPU | S2_219944439 | 2 | 219944439 | 0.666 | $1.13 \times 10^{-9}$ | 0.000176 | yes | chr2:219.94 |
| low-stress | FarmCPU | S2_241935395 | 2 | 241935395 | 0.356 | $1.03 \times 10^{-6}$ | 0.0292 | no | |
| low-stress | BLINK | S3_6763114 | 3 | 6763114 | −0.279 | $2.50 \times 10^{-9}$ | $5.97 \times 10^{-5}$ | yes | chr3:6.76 |
| low-stress | BLINK | S3_35679704 | 3 | 35679704 | −0.597 | $3.81 \times 10^{-10}$ | $1.08 \times 10^{-5}$ | yes | chr3:35.68 |
| low-stress | FarmCPU | S3_124692963 | 3 | 124692963 | −0.291 | $7.51 \times 10^{-8}$ | 0.00292 | yes | chr3:124.69 |
| low-stress | FarmCPU | S3_133835273 | 3 | 133835273 | 0.457 | $5.46 \times 10^{-8}$ | 0.00243 | yes | chr3:133.84 |
| low-stress | BLINK | S3_173688169 | 3 | 173688169 | −0.415 | $5.83 \times 10^{-11}$ | $2.01 \times 10^{-6}$ | yes | chr3:173.69 |
| low-stress | BLINK | S3_180337080 | 3 | 180337080 | 0.283 | $7.61 \times 10^{-8}$ | 0.00139 | yes | chr3:180.34 |
| low-stress | BLINK | S3_180768817 | 3 | 180768817 | −0.208 | $2.23 \times 10^{-6}$ | 0.0248 | no | |
| low-stress | BLINK | S3_220390859 | 3 | 220390859 | −0.712 | $5.59 \times 10^{-14}$ | $5.79 \times 10^{-9}$ | yes | chr3:220.39 |
| low-stress | FarmCPU | S3_220390859 | 3 | 220390859 | −0.487 | $5.53 \times 10^{-9}$ | 0.00043 | yes | chr3:220.39 |
| low-stress | FarmCPU | S5_82949312 | 5 | 82949312 | −0.333 | $1.56 \times 10^{-6}$ | 0.0346 | no | |
| low-stress | BLINK | S6_98464330 | 6 | 98464330 | 0.373 | $5.58 \times 10^{-13}$ | $4.34 \times 10^{-8}$ | yes | chr6:98.46 |
| low-stress | BLINK | S6_133165653 | 6 | 133165653 | −0.494 | $2.18 \times 10^{-12}$ | $9.68 \times 10^{-8}$ | yes | chr6:133.17 |
| low-stress | FarmCPU | S6_133165929 | 6 | 133165929 | −0.616 | $2.11 \times 10^{-8}$ | 0.00131 | yes | chr6:133.17 |
| low-stress | BLINK | S6_174643108 | 6 | 174643108 | −0.437 | $1.71 \times 10^{-6}$ | 0.0197 | no | |

*Supplementary Table S3, continued*

| Analysis | Method | SNP | Chr | Position (bp) | Effect (t/ha) | *P* | FDR (BH) | Bonferroni threshold | Locus (Table S4) |
|---|---|---|---|---|---|---|---|---|---|
| low-stress | BLINK | S7_139494795 | 7 | 139494795 | 0.341 | $6.60 \times 10^{-12}$ | $2.56 \times 10^{-7}$ | yes | chr7:139.49 |
| low-stress | FarmCPU | S7_139494795 | 7 | 139494795 | 0.326 | $3.31 \times 10^{-9}$ | 0.000344 | yes | chr7:139.49 |
| low-stress | FarmCPU | S8_20202171 | 8 | 20202171 | −0.258 | $2.61 \times 10^{-8}$ | 0.00135 | yes | chr8:20.20 |
| low-stress | BLINK | S8_85482596 | 8 | 85482596 | −0.556 | $9.28 \times 10^{-8}$ | 0.00152 | yes | chr8:85.48 |
| low-stress | BLINK | S8_176626806 | 8 | 176626806 | −0.279 | $1.80 \times 10^{-9}$ | $4.66 \times 10^{-5}$ | yes | chr8:176.63 |
| low-stress | BLINK | S9_6864884 | 9 | 6864884 | −0.291 | $5.08 \times 10^{-8}$ | 0.000988 | yes | chr9:6.86 |
| low-stress | BLINK | S9_105639177 | 9 | 105639177 | 0.243 | $5.75 \times 10^{-7}$ | 0.00813 | yes | chr9:105.64 |
| low-stress | FarmCPU | S9_105639177 | 9 | 105639177 | 0.275 | $6.65 \times 10^{-7}$ | 0.023 | no | chr9:105.64 |
| low-stress | BLINK | S9_144221997 | 9 | 144221997 | −0.216 | $1.62 \times 10^{-6}$ | 0.0193 | no | |
| low-stress | BLINK | S10_3534124 | 10 | 3534124 | 0.247 | $1.59 \times 10^{-6}$ | 0.0193 | no | |
| low-stress | FarmCPU | S10_80428278 | 10 | 80428278 | −0.277 | $8.02 \times 10^{-7}$ | 0.0249 | no | |
| low-stress | FarmCPU | S10_148702981 | 10 | 148702981 | −0.223 | $1.30 \times 10^{-6}$ | 0.0321 | no | |
| high-stress | BLINK | S1_259777148 | 1 | 259777148 | 0.351 | $1.75 \times 10^{-8}$ | 0.00077 | yes | chr1:259.78 |
| high-stress | BLINK | S1_264908959 | 1 | 264908959 | −0.198 | $2.20 \times 10^{-6}$ | 0.0402 | no | |
| high-stress | FarmCPU | S1_301883450 | 1 | 301883450 | 0.316 | $6.13 \times 10^{-10}$ | $9.53 \times 10^{-5}$ | yes | chr1:301.88 |
| high-stress | BLINK | S2_131432835 | 2 | 131432835 | −0.256 | $1.47 \times 10^{-8}$ | 0.00077 | yes | chr2:131.43 |
| high-stress | BLINK | S3_108971010 | 3 | 108971010 | −0.522 | $4.26 \times 10^{-10}$ | $6.62 \times 10^{-5}$ | yes | chr3:108.97 |
| high-stress | BLINK | S4_41641600 | 4 | 41641600 | −0.445 | $4.70 \times 10^{-7}$ | 0.0122 | yes | chr4:41.64 |
| high-stress | BLINK | S4_129831466 | 4 | 129831466 | 0.458 | $4.48 \times 10^{-8}$ | 0.00128 | yes | chr4:129.83 |
| high-stress | BLINK | S4_151335090 | 4 | 151335090 | 0.246 | $1.96 \times 10^{-8}$ | 0.00077 | yes | chr4:151.34 |
| high-stress | BLINK | S4_187390116 | 4 | 187390116 | −0.439 | $1.98 \times 10^{-8}$ | 0.00077 | yes | chr4:187.39 |
| high-stress | BLINK | S5_14106884 | 5 | 14106884 | 0.314 | $3.04 \times 10^{-9}$ | 0.000237 | yes | chr5:14.11 |
| high-stress | FarmCPU | S5_14106884 | 5 | 14106884 | 0.313 | $4.06 \times 10^{-8}$ | 0.00421 | yes | chr5:14.11 |
| high-stress | BLINK | S5_19694316 | 5 | 19694316 | −0.254 | $1.14 \times 10^{-6}$ | 0.0222 | no | |
| high-stress | BLINK | S5_19694434 | 5 | 19694434 | −0.254 | $1.14 \times 10^{-6}$ | 0.0222 | no | |
| high-stress | BLINK | S5_145934828 | 5 | 145934828 | 0.466 | $5.98 \times 10^{-7}$ | 0.014 | yes | chr5:145.93 |
| high-stress | FarmCPU | S5_168453024 | 5 | 168453024 | −0.396 | $1.17 \times 10^{-7}$ | 0.00623 | yes | chr5:168.45 |
| high-stress | BLINK | S5_204274986 | 5 | 204274986 | 0.272 | $1.38 \times 10^{-9}$ | 0.000143 | yes | chr5:204.27 |
| high-stress | FarmCPU | S6_88168925 | 6 | 88168925 | −0.387 | $1.20 \times 10^{-7}$ | 0.00623 | yes | chr6:88.17 |
| high-stress | FarmCPU | S6_155998044 | 6 | 155998044 | 0.450 | $4.27 \times 10^{-7}$ | 0.0166 | yes | chr6:156.00 |
| high-stress | FarmCPU | S7_55328932 | 7 | 55328932 | 0.341 | $1.19 \times 10^{-7}$ | 0.00623 | yes | chr7:55.33 |
| high-stress | BLINK | S8_138511916 | 8 | 138511916 | −0.547 | $3.31 \times 10^{-8}$ | 0.00114 | yes | chr8:138.51 |
| high-stress | BLINK | S8_167941186 | 8 | 167941186 | −0.330 | $4.52 \times 10^{-8}$ | 0.00128 | yes | chr8:167.94 |

*Supplementary Table S3, continued*

| Analysis | Method | SNP | Chr | Position (bp) | Effect (t/ha) | *P* | FDR (BH) | Bonferroni threshold | Locus (Table S4) |
|---|---|---|---|---|---|---|---|---|---|
| high-stress | FarmCPU | S8_171403871 | 8 | 171403871 | 0.547 | $1.68 \times 10^{-10}$ | $5.24 \times 10^{-5}$ | yes | chr8:171.40 |
| high-stress | BLINK | S8_171406449 | 8 | 171406449 | 0.419 | $4.04 \times 10^{-10}$ | $6.62 \times 10^{-5}$ | yes | chr8:171.40 |
| high-stress | FarmCPU | S8_171406449 | 8 | 171406449 | 0.351 | $2.38 \times 10^{-7}$ | 0.0106 | yes | chr8:171.40 |
| high-stress | BLINK | S10_143856591 | 10 | 143856591 | −0.219 | $6.30 \times 10^{-7}$ | 0.014 | no | |
| delta (low − high) | BLINK | S1_193836307 | 1 | 193836307 | −0.469 | $1.20 \times 10^{-11}$ | $1.25 \times 10^{-6}$ | yes | chr1:193.84 |
| delta (low − high) | BLINK | S1_206313491 | 1 | 206313491 | 0.820 | $3.15 \times 10^{-11}$ | $2.45 \times 10^{-6}$ | yes | chr1:206.31 |
| delta (low − high) | BLINK | S1_307021453 | 1 | 307021453 | −0.259 | $2.98 \times 10^{-7}$ | 0.00772 | yes | chr1:307.02 |
| delta (low − high) | BLINK | S2_219316588 | 2 | 219316588 | 0.392 | $8.58 \times 10^{-8}$ | 0.00267 | yes | chr2:219.32 |
| delta (low − high) | BLINK | S2_230061580 | 2 | 230061580 | 0.389 | $4.82 \times 10^{-10}$ | $2.50 \times 10^{-5}$ | yes | chr2:230.06 |
| delta (low − high) | FarmCPU | S2_241776807 | 2 | 241776807 | −0.510 | $1.54 \times 10^{-7}$ | 0.024 | yes | chr2:241.78 |
| delta (low − high) | BLINK | S3_9272116 | 3 | 9272116 | −0.412 | $9.15 \times 10^{-12}$ | $1.25 \times 10^{-6}$ | yes | chr3:9.27 |
| delta (low − high) | BLINK | S3_180336777 | 3 | 180336777 | −0.335 | $4.78 \times 10^{-7}$ | 0.0114 | yes | chr3:180.34 |
| delta (low − high) | FarmCPU | S4_129831941 | 4 | 129831941 | −0.351 | $1.00 \times 10^{-7}$ | 0.024 | yes | chr4:129.83 |
| delta (low − high) | BLINK | S5_4712053 | 5 | 4712053 | −0.594 | $2.33 \times 10^{-10}$ | $1.45 \times 10^{-5}$ | yes | chr5:4.71 |
| delta (low − high) | BLINK | S5_98789580 | 5 | 98789580 | −0.371 | $1.75 \times 10^{-7}$ | 0.00494 | yes | chr5:98.79 |
| delta (low − high) | FarmCPU | S5_163132214 | 5 | 163132214 | −0.354 | $3.80 \times 10^{-7}$ | 0.0394 | yes | chr5:163.13 |
| delta (low − high) | BLINK | S5_170949929 | 5 | 170949929 | −0.519 | $5.15 \times 10^{-9}$ | 0.000229 | yes | chr5:170.95 |
| delta (low − high) | BLINK | S6_33313575 | 6 | 33313575 | 0.372 | $9.63 \times 10^{-7}$ | 0.0187 | no | |
| delta (low − high) | BLINK | S6_166672717 | 6 | 166672717 | −0.439 | $2.53 \times 10^{-8}$ | 0.000875 | yes | chr6:166.67 |
| delta (low − high) | BLINK | S7_1299390 | 7 | 1299390 | −0.266 | $6.87 \times 10^{-7}$ | 0.0149 | no | |
| delta (low − high) | BLINK | S8_7116806 | 8 | 7116806 | 0.312 | $2.16 \times 10^{-8}$ | 0.000838 | yes | chr8:7.12 |
| delta (low − high) | BLINK | S8_15293321 | 8 | 15293321 | 0.555 | $2.49 \times 10^{-18}$ | $7.75 \times 10^{-13}$ | yes | chr8:15.29 |
| delta (low − high) | BLINK | S10_4117886 | 10 | 4117886 | −0.272 | $7.20 \times 10^{-7}$ | 0.0149 | no | |

**Supplementary Table S4:** Loci associated with grain yield under low stress, under high stress and in the delta (low − high stress), named by the position (chromosome and Mb) of the lead SNP. The *No.* column gives the number of the locus in Figure 7 or, for the delta, in Supplementary Figure S4. The *Method* column indicates the method with which the locus reached the Bonferroni threshold (section 2.11), and *P* is that of the lead SNP in the analysis in which the locus was identified. MAF is that of the lead SNP in the parents of the model. PVE is the squared effect of the lead SNP times the variance of its genotype, divided by the phenotypic variance, in the hybrids of the analysis in which the locus was identified. It tends to be overestimated. *Effect* is the effect of the favorable allele of the lead SNP, in t/ha, in the three analyses. For SNPs with low MAF, it also tends to be overestimated. The effect for delta is the one estimated in the GWAS of the delta, with the 2,084 hybrids evaluated in both conditions, not the difference between the low- and high-stress effects. *SD ratio high/low* is the standard deviation of the local GEBV under high stress divided by that under low stress, among the parents of the model, in the 164-kb segment that contains the lead SNP. Across the genome, the median of this ratio is 0.75. The upper third, the segments that lose the least variance under stress, is above 0.83, and the lower third is below 0.68. The last column indicates, for each heterotic group (Stiff Stalk / non-Stiff Stalk / Iodent), whether the 164-kb segment of the lead SNP is a high-effect, high-variance segment in the condition of the locus (section 2.11; Figure 7b), and does not apply to the delta loci. Two delta loci lie less than 1 kb from the lead SNP of a low- or high-stress locus: chr3:180.34 (D8 and L16) and chr4:129.83 (D9 and H6).

| Analysis | *No.* | Locus | *Method* | *P* (lead SNP) | MAF | PVE (%) | *Effect*, low-stress (t/ha) | *Effect*, high-stress (t/ha) | *Effect*, delta (t/ha) | *SD ratio high/low*, 164 kb | High-effect, high-variance segment (Stiff Stalk / non-Stiff Stalk / Iodent) |
|---|---|---|---|---|---|---|---|---|---|---|---|
| Low-stress | L1 | chr1:7.92 | BLINK | $7.6 \times 10^{-11}$ | 0.11 | 2.4 | 0.47 | 0.08 | 0.07 | 0.69 | yes / no / yes |
| Low-stress | L2 | chr1:11.49 | BLINK | $1.2 \times 10^{-14}$ | 0.47 | 4.8 | 0.45 | 0.09 | 0.17 | 0.73 | yes / yes / yes |
| Low-stress | L3 | chr1:19.87 | BLINK | $2.0 \times 10^{-12}$ | 0.09 | 1.9 | 0.47 | 0.10 | 0.13 | 0.92 | yes / yes / yes |
| Low-stress | L4 | chr1:209.41 | BLINK | $1.2 \times 10^{-12}$ | 0.08 | 2.0 | 0.56 | 0.11 | 0.17 | 1.03 | yes / yes / yes |
| Low-stress | L5 | chr1:218.12 | BLINK | $1.5 \times 10^{-17}$ | 0.08 | 3.8 | 0.65 | 0.21 | −0.01 | 0.88 | no / no / yes |
| Low-stress | L6 | chr2:0.88 | BLINK | $9.7 \times 10^{-9}$ | 0.12 | 1.9 | 0.36 | 0.05 | 0.15 | 0.94 | no / no / no |
| Low-stress | L7 | chr2:13.82 | BLINK | $3.5 \times 10^{-7}$ | 0.26 | 0.6 | 0.23 | 0.05 | 0.05 | 1.07 | yes / yes / yes |
| Low-stress | L8 | chr2:29.66 | BLINK/FarmCPU | $7.9 \times 10^{-10}$ | 0.18 | 2.4 | 0.39 | 0.14 | 0.02 | 0.76 | yes / no / yes |
| Low-stress | L9 | chr2:201.07 | BLINK | $8.2 \times 10^{-8}$ | 0.24 | 1.0 | 0.24 | 0.14 | −0.01 | 0.87 | yes / yes / yes |
| Low-stress | L10 | chr2:219.94 | BLINK/FarmCPU | $1.1 \times 10^{-9}$ | 0.05 | 1.0 | 0.67 | −0.07 | 0.39 | 0.58 | no / no / yes |
| Low-stress | L11 | chr3:6.76 | BLINK | $2.5 \times 10^{-9}$ | 0.24 | 1.0 | 0.28 | 0.13 | 0.13 | 0.79 | yes / yes / yes |
| Low-stress | L12 | chr3:35.68 | BLINK | $3.8 \times 10^{-10}$ | 0.06 | 1.2 | 0.60 | 0.19 | −0.03 | 0.62 | yes / yes / yes |
| Low-stress | L13 | chr3:124.69 | FarmCPU | $7.5 \times 10^{-8}$ | 0.39 | 1.7 | 0.29 | 0.05 | −0.08 | 0.64 | yes / yes / yes |
| Low-stress | L14 | chr3:133.84 | FarmCPU | $5.5 \times 10^{-8}$ | 0.08 | 2.2 | 0.46 | 0.07 | 0.26 | 0.53 | no / no / no |
| Low-stress | L15 | chr3:173.69 | BLINK | $5.8 \times 10^{-11}$ | 0.11 | 2.0 | 0.42 | 0.03 | −0.09 | 0.44 | yes / yes / no |
| Low-stress | L16 | chr3:180.34 | BLINK | $7.6 \times 10^{-8}$ | 0.18 | 1.3 | 0.28 | 0.05 | 0.12 | 0.45 | yes / no / no |
| Low-stress | L17 | chr3:220.39 | BLINK/FarmCPU | $5.6 \times 10^{-14}$ | 0.09 | 2.5 | 0.71 | 0.16 | 0.18 | 0.69 | yes / yes / yes |
| Low-stress | L18 | chr6:98.46 | BLINK | $5.6 \times 10^{-13}$ | 0.46 | 2.9 | 0.37 | 0.12 | −0.07 | 0.90 | yes / yes / yes |
| Low-stress | L19 | chr6:133.17 | BLINK/FarmCPU | $2.2 \times 10^{-12}$ | 0.09 | 2.1 | 0.49 | 0.15 | 0.05 | 0.56 | no / no / yes |
| Low-stress | L20 | chr7:139.49 | BLINK/FarmCPU | $6.6 \times 10^{-12}$ | 0.26 | 2.0 | 0.34 | 0.09 | 0.12 | 0.75 | yes / yes / yes |
| Low-stress | L21 | chr8:20.20 | FarmCPU | $2.6 \times 10^{-8}$ | 0.39 | 1.2 | 0.26 | 0.06 | 0.09 | 0.76 | yes / yes / yes |
| Low-stress | L22 | chr8:85.48 | BLINK | $9.3 \times 10^{-8}$ | 0.05 | 0.8 | 0.56 | 0.30 | 0.19 | 0.64 | no / yes / no |

*Supplementary Table S4, continued*

| Analysis | *No.* | Locus | *Method* | *P* (lead SNP) | MAF | PVE (%) | *Effect*, low-stress (t/ha) | *Effect*, high-stress (t/ha) | *Effect*, delta (t/ha) | *SD ratio high/low*, 164 kb | High-effect, high-variance segment (Stiff Stalk / non-Stiff Stalk / Iodent) |
|---|---|---|---|---|---|---|---|---|---|---|---|
| Low-stress | L23 | chr8:176.63 | BLINK | $1.8 \times 10^{-9}$ | 0.39 | 1.3 | 0.28 | 0.13 | 0.10 | 0.79 | yes / no / no |
| Low-stress | L24 | chr9:6.86 | BLINK | $5.1 \times 10^{-8}$ | 0.39 | 1.3 | 0.29 | −0.04 | 0.15 | 0.84 | no / no / no |
| Low-stress | L25 | chr9:105.64 | BLINK | $5.8 \times 10^{-7}$ | 0.20 | 0.9 | 0.24 | 0.12 | 0.10 | 0.73 | no / yes / yes |
| High-stress | H1 | chr1:259.78 | BLINK | $1.7 \times 10^{-8}$ | 0.28 | 1.7 | 0.11 | 0.35 | 0.03 | 0.68 | yes / no / yes |
| High-stress | H2 | chr1:301.88 | FarmCPU | $6.1 \times 10^{-10}$ | 0.41 | 2.8 | 0.10 | 0.32 | −0.09 | 0.78 | yes / yes / yes |
| High-stress | H3 | chr2:131.43 | BLINK | $1.5 \times 10^{-8}$ | 0.28 | 1.3 | 0.04 | 0.26 | −0.13 | 0.69 | no / no / no |
| High-stress | H4 | chr3:108.97 | BLINK | $4.3 \times 10^{-10}$ | 0.06 | 1.1 | 0.30 | 0.52 | −0.29 | 0.67 | no / no / no |
| High-stress | H5 | chr4:41.64 | BLINK | $4.7 \times 10^{-7}$ | 0.07 | 0.8 | −0.05 | 0.45 | −0.14 | 0.74 | no / yes / yes |
| High-stress | H6 | chr4:129.83 | BLINK | $4.5 \times 10^{-8}$ | 0.09 | 1.5 | 0.10 | 0.46 | −0.15 | 1.31 | no / no / no |
| High-stress | H7 | chr4:151.34 | BLINK | $2.0 \times 10^{-8}$ | 0.36 | 1.4 | 0.04 | 0.25 | −0.08 | 0.85 | yes / yes / yes |
| High-stress | H8 | chr4:187.39 | BLINK | $2.0 \times 10^{-8}$ | 0.07 | 1.3 | 0.09 | 0.44 | −0.03 | 0.79 | yes / yes / no |
| High-stress | H9 | chr5:14.11 | BLINK/FarmCPU | $3.0 \times 10^{-9}$ | 0.20 | 1.3 | 0.06 | 0.31 | 0.10 | 0.99 | yes / yes / yes |
| High-stress | H10 | chr5:145.93 | BLINK | $6.0 \times 10^{-7}$ | 0.06 | 0.6 | 0.08 | 0.47 | −0.20 | 1.16 | no / no / no |
| High-stress | H11 | chr5:168.45 | FarmCPU | $1.2 \times 10^{-7}$ | 0.36 | 2.2 | 0.18 | 0.40 | −0.07 | 0.64 | no / no / no |
| High-stress | H12 | chr5:204.27 | BLINK | $1.4 \times 10^{-9}$ | 0.49 | 1.4 | −0.07 | 0.27 | −0.19 | 0.60 | no / no / no |
| High-stress | H13 | chr6:88.17 | FarmCPU | $1.2 \times 10^{-7}$ | 0.30 | 2.3 | 0.15 | 0.39 | 0.05 | 1.06 | no / no / no |
| High-stress | H14 | chr6:156.00 | FarmCPU | $4.3 \times 10^{-7}$ | 0.31 | 3.1 | 0.02 | 0.45 | −0.13 | 0.93 | yes / yes / yes |
| High-stress | H15 | chr7:55.33 | FarmCPU | $1.2 \times 10^{-7}$ | 0.27 | 1.3 | −0.02 | 0.34 | −0.13 | 0.78 | no / no / no |
| High-stress | H16 | chr8:138.51 | BLINK | $3.3 \times 10^{-8}$ | 0.06 | 0.9 | −0.16 | 0.55 | −0.40 | 0.69 | yes / yes / yes |
| High-stress | H17 | chr8:167.94 | BLINK | $4.5 \times 10^{-8}$ | 0.20 | 0.8 | 0.07 | 0.33 | 0.04 | 0.84 | yes / yes / yes |
| High-stress | H18 | chr8:171.40 | BLINK/FarmCPU | $1.7 \times 10^{-10}$ | 0.12 | 2.2 | 0.02 | 0.55 | −0.15 | 0.76 | yes / yes / yes |
| Delta (low − high) | D1 | chr1:193.84 | BLINK | $1.2 \times 10^{-11}$ | 0.25 | 3.7 | 0.16 | −0.06 | 0.47 | 0.69 | – |
| Delta (low − high) | D2 | chr1:206.31 | BLINK | $3.2 \times 10^{-11}$ | 0.06 | 1.5 | 0.09 | −0.08 | 0.82 | 0.60 | – |
| Delta (low − high) | D3 | chr1:307.02 | BLINK | $3.0 \times 10^{-7}$ | 0.37 | 0.8 | 0.07 | −0.06 | 0.26 | 0.73 | – |
| Delta (low − high) | D4 | chr2:219.32 | BLINK | $8.6 \times 10^{-8}$ | 0.12 | 1.6 | 0.14 | 0.03 | 0.39 | 0.71 | – |
| Delta (low − high) | D5 | chr2:230.06 | BLINK | $4.8 \times 10^{-10}$ | 0.18 | 1.8 | 0.08 | −0.12 | 0.39 | 0.69 | – |
| Delta (low − high) | D6 | chr2:241.78 | FarmCPU | $1.5 \times 10^{-7}$ | 0.11 | 2.9 | 0.22 | −0.20 | 0.51 | 0.78 | – |
| Delta (low − high) | D7 | chr3:9.27 | BLINK | $9.2 \times 10^{-12}$ | 0.26 | 2.1 | 0.09 | −0.04 | 0.41 | 0.65 | – |
| Delta (low − high) | D8 | chr3:180.34 | BLINK | $4.8 \times 10^{-7}$ | 0.23 | 0.9 | 0.16 | −0.05 | 0.33 | 0.45 | – |
| Delta (low − high) | D9 | chr4:129.83 | FarmCPU | $1.0 \times 10^{-7}$ | 0.36 | 2.3 | −0.03 | −0.02 | 0.35 | 1.31 | – |
| Delta (low − high) | D10 | chr5:4.71 | BLINK | $2.3 \times 10^{-10}$ | 0.06 | 1.7 | 0.20 | −0.06 | 0.59 | 0.79 | – |
| Delta (low − high) | D11 | chr5:98.79 | BLINK | $1.7 \times 10^{-7}$ | 0.12 | 1.1 | 0.02 | −0.05 | 0.37 | 0.61 | – |
| Delta (low − high) | D12 | chr5:163.13 | FarmCPU | $3.8 \times 10^{-7}$ | 0.34 | 1.7 | 0.15 | −0.08 | 0.35 | 0.64 | – |
| Delta (low − high) | D13 | chr5:170.95 | BLINK | $5.2 \times 10^{-9}$ | 0.14 | 1.1 | 0.16 | −0.14 | 0.52 | 0.77 | – |
| Delta (low − high) | D14 | chr6:166.67 | BLINK | $2.5 \times 10^{-8}$ | 0.06 | 1.1 | 0.03 | −0.20 | 0.44 | 1.05 | – |
| Delta (low − high) | D15 | chr8:7.12 | BLINK | $2.2 \times 10^{-8}$ | 0.46 | 1.6 | 0.05 | −0.08 | 0.31 | 0.56 | – |

*Supplementary Table S4, continued*

| Analysis | *No.* | Locus | *Method* | *P* (lead SNP) | MAF | PVE (%) | *Effect*, low-stress (t/ha) | *Effect*, high-stress (t/ha) | *Effect*, delta (t/ha) | *SD ratio high/low*, 164 kb | High-effect, high-variance segment (Stiff Stalk / non-Stiff Stalk / Iodent) |
|---|---|---|---|---|---|---|---|---|---|---|---|
| Delta (low − high) | D16 | chr8:15.29 | BLINK | $2.5 \times 10^{-18}$ | 0.21 | 4.5 | −0.02 | −0.05 | 0.56 | 0.75 | – |

**Supplementary Table S5:** Correlation of the local GEBV between the two conditions ($r_j$), segment by segment, in each heterotic group. $r_j$ of each 164-kb segment is calculated among the parents of the group in the model. For each group, the table gives the number of segments and the median $r_j$. It also gives how many segments have $r_j$ below 0.8 and the percentage of the segment variance (Figure 4) that they account for under low and under high stress. The last column shows how many of the segments in the top 20% for variance in both conditions have $r_j$ below 0.8.

| Group | Segments | Median $r_j$ | Segments with $r_j < 0.8$ | Share of the low-stress segment variance with $r_j < 0.8$ (%) | Share of the high-stress segment variance with $r_j < 0.8$ (%) | Top 20% in both conditions | Top 20% with $r_j < 0.8$ |
|---|---|---|---|---|---|---|---|
| Stiff Stalk | 10044 | 0.99 | 1363 (14%) | 2.4 | 2.1 | 1658 | 13 (0.8%) |
| non-Stiff Stalk | 10045 | 0.96 | 2269 (23%) | 6.8 | 7.2 | 1492 | 32 (2.1%) |
| Iodent | 10017 | 1.00 | 54 (0.5%) | 0.0 | 0.0 | 1958 | 0 (0.0%) |

**Supplementary Table S6:** Sets of 2–50 founders by pool, condition and rule (stacking and truncation). The table gives the potential, in t/ha above the mean of the parents of the model and as a percentage of the potential of the whole pool, the mean GEBV, the mean coancestry among the founders and over all pairs of lines in the pool, the different versions per 164-kb segment, the founders with the most common version and the expected heterozygosity. The versions and the coancestry of the pool were calculated only for 8 or more founders.

| Pool | Condition | Founders (n) | Rule | Potential of the founders (t/ha) | Potential, % of the whole pool | Potential of the whole pool (t/ha) | Mean GEBV of the founders (t/ha) | Mean coancestry among founders | Mean coancestry, average pair of the pool | Versions per 164-kb segment | Founders sharing the most common version | Expected heterozygosity |
|---|---|---|---|---|---|---|---|---|---|---|---|---|
| Stiff Stalk pool | low-stress | 2 | stacking | 1.71 | 17.5 | 9.80 | 1.71 | 0.221 | – | – | – | 0.136 |
| Stiff Stalk pool | low-stress | 2 | truncation | 1.71 | 17.5 | 9.80 | 1.71 | 0.221 | – | – | – | 0.136 |
| Stiff Stalk pool | high-stress | 2 | stacking | 1.25 | 17.4 | 7.21 | 1.25 | 0.077 | – | – | – | 0.168 |
| Stiff Stalk pool | high-stress | 2 | truncation | 1.25 | 17.4 | 7.21 | 1.25 | 0.077 | – | – | – | 0.168 |
| Stiff Stalk pool | low-stress | 4 | stacking | 5.86 | 59.8 | 9.80 | 1.40 | 0.050 | – | – | – | 0.258 |
| Stiff Stalk pool | low-stress | 4 | truncation | 5.53 | 56.5 | 9.80 | 1.52 | 0.128 | – | – | – | 0.228 |
| Stiff Stalk pool | high-stress | 4 | stacking | 4.38 | 60.8 | 7.21 | 1.11 | 0.036 | – | – | – | 0.265 |
| Stiff Stalk pool | high-stress | 4 | truncation | 4.38 | 60.8 | 7.21 | 1.11 | 0.036 | – | – | – | 0.265 |
| Stiff Stalk pool | low-stress | 6 | stacking | 7.28 | 74.3 | 9.80 | 0.96 | 0.030 | – | – | – | 0.296 |
| Stiff Stalk pool | low-stress | 6 | truncation | 6.88 | 70.2 | 9.80 | 1.43 | 0.128 | – | – | – | 0.256 |
| Stiff Stalk pool | high-stress | 6 | stacking | 5.45 | 75.6 | 7.21 | 0.92 | 0.042 | – | – | – | 0.291 |
| Stiff Stalk pool | high-stress | 6 | truncation | 5.03 | 69.8 | 7.21 | 1.03 | 0.156 | – | – | – | 0.261 |
| Stiff Stalk pool | low-stress | 8 | stacking | 8.06 | 82.3 | 9.80 | 0.96 | 0.040 | 0.154 | 4.3 | 3.8 | 0.308 |
| Stiff Stalk pool | low-stress | 8 | truncation | 7.49 | 76.4 | 9.80 | 1.38 | 0.145 | 0.154 | 3.5 | 4.5 | 0.265 |
| Stiff Stalk pool | high-stress | 8 | stacking | 5.94 | 82.4 | 7.21 | 0.68 | 0.045 | 0.154 | 4.4 | 3.8 | 0.308 |
| Stiff Stalk pool | high-stress | 8 | truncation | 5.21 | 72.3 | 7.21 | 0.98 | 0.229 | 0.154 | 3.3 | 4.6 | 0.251 |
| Stiff Stalk pool | low-stress | 12 | stacking | 8.71 | 88.9 | 9.80 | 0.62 | 0.052 | 0.154 | 5.3 | 5.4 | 0.322 |
| Stiff Stalk pool | low-stress | 12 | truncation | 8.18 | 83.5 | 9.80 | 1.28 | 0.161 | 0.154 | 4.3 | 6.8 | 0.270 |
| Stiff Stalk pool | high-stress | 12 | stacking | 6.43 | 89.3 | 7.21 | 0.55 | 0.044 | 0.154 | 5.4 | 5.4 | 0.323 |
| Stiff Stalk pool | high-stress | 12 | truncation | 5.90 | 81.9 | 7.21 | 0.92 | 0.192 | 0.154 | 4.2 | 6.6 | 0.271 |
| Stiff Stalk pool | low-stress | 16 | stacking | 8.99 | 91.8 | 9.80 | 0.51 | 0.050 | 0.154 | 6.0 | 7.1 | 0.330 |
| Stiff Stalk pool | low-stress | 16 | truncation | 8.50 | 86.8 | 9.80 | 1.22 | 0.175 | 0.154 | 4.8 | 9.1 | 0.272 |
| Stiff Stalk pool | high-stress | 16 | stacking | 6.64 | 92.1 | 7.21 | 0.43 | 0.052 | 0.154 | 6.0 | 7.1 | 0.329 |
| Stiff Stalk pool | high-stress | 16 | truncation | 6.08 | 84.4 | 7.21 | 0.86 | 0.210 | 0.154 | 4.6 | 8.9 | 0.269 |
| Stiff Stalk pool | low-stress | 20 | stacking | 9.15 | 93.4 | 9.80 | 0.39 | 0.062 | 0.154 | 6.8 | 9.0 | 0.327 |
| Stiff Stalk pool | low-stress | 20 | truncation | 8.67 | 88.4 | 9.80 | 1.17 | 0.177 | 0.154 | 5.1 | 11.2 | 0.274 |
| Stiff Stalk pool | high-stress | 20 | stacking | 6.74 | 93.5 | 7.21 | 0.25 | 0.050 | 0.154 | 6.8 | 8.9 | 0.333 |
| Stiff Stalk pool | high-stress | 20 | truncation | 6.32 | 87.7 | 7.21 | 0.81 | 0.205 | 0.154 | 5.1 | 11.4 | 0.270 |
| Stiff Stalk pool | low-stress | 25 | stacking | 9.27 | 94.6 | 9.80 | 0.38 | 0.061 | 0.154 | 7.6 | 11.4 | 0.328 |
| Stiff Stalk pool | low-stress | 25 | truncation | 8.80 | 89.8 | 9.80 | 1.12 | 0.188 | 0.154 | 5.5 | 14.2 | 0.274 |

*Supplementary Table S6, continued*

| Pool | Condition | Founders (n) | Rule | Potential of the founders (t/ha) | Potential, % of the whole pool | Potential of the whole pool (t/ha) | Mean GEBV of the founders (t/ha) | Mean coancestry among founders | Mean coancestry, average pair of the pool | Versions per 164-kb segment | Founders sharing the most common version | Expected heterozygosity |
|---|---|---|---|---|---|---|---|---|---|---|---|---|
| Stiff Stalk pool | high-stress | 25 | stacking | 6.84 | 94.9 | 7.21 | 0.25 | 0.057 | 0.154 | 7.8 | 10.9 | 0.335 |
| Stiff Stalk pool | high-stress | 25 | truncation | 6.45 | 89.5 | 7.21 | 0.76 | 0.211 | 0.154 | 5.5 | 14.5 | 0.270 |
| Stiff Stalk pool | low-stress | 50 | stacking | 9.52 | 97.1 | 9.80 | 0.16 | 0.084 | 0.154 | 10.9 | 23.5 | 0.325 |
| Stiff Stalk pool | low-stress | 50 | truncation | 9.08 | 92.6 | 9.80 | 0.92 | 0.188 | 0.154 | 6.7 | 27.8 | 0.281 |
| Stiff Stalk pool | high-stress | 50 | stacking | 7.01 | 97.4 | 7.21 | 0.15 | 0.081 | 0.154 | 10.9 | 23.1 | 0.327 |
| Stiff Stalk pool | high-stress | 50 | truncation | 6.81 | 94.5 | 7.21 | 0.60 | 0.202 | 0.154 | 7.4 | 28.9 | 0.278 |
| non-Stiff Stalk pool | low-stress | 2 | stacking | 2.23 | 19.3 | 11.51 | 2.23 | 0.098 | – | – | – | 0.176 |
| non-Stiff Stalk pool | low-stress | 2 | truncation | 2.23 | 19.3 | 11.51 | 2.23 | 0.098 | – | – | – | 0.176 |
| non-Stiff Stalk pool | high-stress | 2 | stacking | 1.96 | 24.4 | 8.03 | 1.96 | 0.462 | – | – | – | 0.084 |
| non-Stiff Stalk pool | high-stress | 2 | truncation | 1.96 | 24.4 | 8.03 | 1.96 | 0.462 | – | – | – | 0.084 |
| non-Stiff Stalk pool | low-stress | 4 | stacking | 6.71 | 58.3 | 11.51 | 2.03 | 0.049 | – | – | – | 0.277 |
| non-Stiff Stalk pool | low-stress | 4 | truncation | 6.65 | 57.8 | 11.51 | 2.09 | 0.072 | – | – | – | 0.276 |
| non-Stiff Stalk pool | high-stress | 4 | stacking | 4.58 | 57.0 | 8.03 | 1.45 | 0.025 | – | – | – | 0.264 |
| non-Stiff Stalk pool | high-stress | 4 | truncation | 3.07 | 38.2 | 8.03 | 1.88 | 0.465 | – | – | – | 0.123 |
| non-Stiff Stalk pool | low-stress | 6 | stacking | 8.30 | 72.1 | 11.51 | 1.84 | 0.045 | – | – | – | 0.313 |
| non-Stiff Stalk pool | low-stress | 6 | truncation | 8.11 | 70.5 | 11.51 | 2.00 | 0.085 | – | – | – | 0.299 |
| non-Stiff Stalk pool | high-stress | 6 | stacking | 5.71 | 71.1 | 8.03 | 1.30 | 0.039 | – | – | – | 0.299 |
| non-Stiff Stalk pool | high-stress | 6 | truncation | 3.56 | 44.3 | 8.03 | 1.83 | 0.434 | – | – | – | 0.149 |
| non-Stiff Stalk pool | low-stress | 8 | stacking | 9.17 | 79.7 | 11.51 | 1.77 | 0.030 | 0.089 | 5.0 | 3.3 | 0.324 |
| non-Stiff Stalk pool | low-stress | 8 | truncation | 8.83 | 76.7 | 11.51 | 1.95 | 0.083 | 0.089 | 4.8 | 3.5 | 0.309 |
| non-Stiff Stalk pool | high-stress | 8 | stacking | 6.28 | 78.2 | 8.03 | 1.18 | 0.032 | 0.089 | 4.6 | 3.5 | 0.316 |
| non-Stiff Stalk pool | high-stress | 8 | truncation | 3.62 | 45.1 | 8.03 | 1.79 | 0.451 | 0.089 | 2.1 | 5.8 | 0.152 |
| non-Stiff Stalk pool | low-stress | 12 | stacking | 10.01 | 87.0 | 11.51 | 1.54 | 0.046 | 0.089 | 6.3 | 4.7 | 0.338 |
| non-Stiff Stalk pool | low-stress | 12 | truncation | 9.68 | 84.1 | 11.51 | 1.83 | 0.076 | 0.089 | 5.9 | 5.1 | 0.322 |
| non-Stiff Stalk pool | high-stress | 12 | stacking | 6.88 | 85.7 | 8.03 | 1.21 | 0.036 | 0.089 | 5.5 | 5.3 | 0.323 |
| non-Stiff Stalk pool | high-stress | 12 | truncation | 4.03 | 50.2 | 8.03 | 1.72 | 0.395 | 0.089 | 2.5 | 8.2 | 0.179 |
| non-Stiff Stalk pool | low-stress | 16 | stacking | 10.41 | 90.4 | 11.51 | 1.46 | 0.044 | 0.089 | 7.3 | 6.2 | 0.345 |
| non-Stiff Stalk pool | low-stress | 16 | truncation | 10.19 | 88.6 | 11.51 | 1.73 | 0.065 | 0.089 | 7.0 | 6.6 | 0.332 |
| non-Stiff Stalk pool | high-stress | 16 | stacking | 7.19 | 89.6 | 8.03 | 0.99 | 0.031 | 0.089 | 6.7 | 6.7 | 0.336 |
| non-Stiff Stalk pool | high-stress | 16 | truncation | 4.04 | 50.4 | 8.03 | 1.68 | 0.407 | 0.089 | 2.6 | 11.0 | 0.178 |
| non-Stiff Stalk pool | low-stress | 20 | stacking | 10.66 | 92.6 | 11.51 | 1.29 | 0.034 | 0.089 | 8.9 | 7.1 | 0.355 |
| non-Stiff Stalk pool | low-stress | 20 | truncation | 10.32 | 89.7 | 11.51 | 1.65 | 0.084 | 0.089 | 7.4 | 8.1 | 0.334 |
| non-Stiff Stalk pool | high-stress | 20 | stacking | 7.40 | 92.2 | 8.03 | 0.93 | 0.033 | 0.089 | 7.6 | 8.0 | 0.343 |
| non-Stiff Stalk pool | high-stress | 20 | truncation | 4.07 | 50.8 | 8.03 | 1.64 | 0.406 | 0.089 | 2.7 | 13.6 | 0.181 |

*Supplementary Table S6, continued*

| Pool | Condition | Founders (n) | Rule | Potential of the founders (t/ha) | Potential, % of the whole pool | Potential of the whole pool (t/ha) | Mean GEBV of the founders (t/ha) | Mean coancestry among founders | Mean coancestry, average pair of the pool | Versions per 164-kb segment | Founders sharing the most common version | Expected heterozygosity |
|---|---|---|---|---|---|---|---|---|---|---|---|---|
| non-Stiff Stalk pool | low-stress | 25 | stacking | 10.84 | 94.2 | 11.51 | 1.17 | 0.036 | 0.089 | 10.0 | 9.0 | 0.357 |
| non-Stiff Stalk pool | low-stress | 25 | truncation | 10.57 | 91.8 | 11.51 | 1.57 | 0.061 | 0.089 | 8.5 | 9.8 | 0.343 |
| non-Stiff Stalk pool | high-stress | 25 | stacking | 7.53 | 93.9 | 8.03 | 0.77 | 0.030 | 0.089 | 8.4 | 9.9 | 0.346 |
| non-Stiff Stalk pool | high-stress | 25 | truncation | 4.46 | 55.6 | 8.03 | 1.60 | 0.376 | 0.089 | 2.9 | 16.6 | 0.193 |
| non-Stiff Stalk pool | low-stress | 50 | stacking | 11.20 | 97.3 | 11.51 | 0.78 | 0.040 | 0.089 | 14.8 | 17.4 | 0.362 |
| non-Stiff Stalk pool | low-stress | 50 | truncation | 11.02 | 95.7 | 11.51 | 1.32 | 0.052 | 0.089 | 11.2 | 19.8 | 0.345 |
| non-Stiff Stalk pool | high-stress | 50 | stacking | 7.81 | 97.2 | 8.03 | 0.39 | 0.031 | 0.089 | 13.2 | 18.4 | 0.357 |
| non-Stiff Stalk pool | high-stress | 50 | truncation | 5.72 | 71.3 | 8.03 | 1.45 | 0.320 | 0.089 | 4.2 | 31.7 | 0.217 |

**Supplementary File S1.** Group and first three principal components of each of the 2,075 parents (Figure 3). File `File_S1.xlsx`.
